# Decoupled genetic and epigenetic variation in the montane endemic *Erodium cazorlanum* (Geraniaceae) and a widespread congener

**DOI:** 10.1101/2024.04.18.589719

**Authors:** Rubén Martín-Blázquez, Mónica Medrano, Pilar Bazaga, Francisco Balao, Ovidiu Paun, Conchita Alonso

## Abstract

Epigenetic states offer an additional layer of variation besides genetic polymorphism that contribute to phenotypic variation and may arise either randomly or in response to environmental factors. We hypothesize that closely related species with different life-histories and habitat requirements could show distinct patterns of intraspecific epigenetic variation. We used Restriction-site Associated DNA sequencing (RADseq) and its bisulfite-converted variant (bsRADseq) to investigate in Sierra de Cazorla (SE Spain) the genetic and epigenetic population structure of two congeneric species, the endemic woody perennial *Erodium cazorlanum* and the widely distributed annual herb *E. cicutarium*. Population genomics analyses revealed no structure in either *E. cazorlanum* and *E. cicutarium,* suggesting substantial gene flow between the study populations. In contrast, we found that the mean proportion of global DNA methylation was different between populations and species, with *E. cazorlanum* DNA showing higher methylation average and across-individuals variation. For each species, we searched for the loci with the strongest epigenetic differentiation between populations (differentially methylated cytosines, DMCs), and summarized them across regions (differentially methylated regions, DMRs). Multivariate analysis and hierarchical clustering of single cytosines’ methylation percentages did not group individuals by population, pointing to high epigenetic variation within populations. DMRs were enriched in *Copia* transposable elements, putatively associated with stress response in plants. Our results suggest that variation at both genetic and epigenetic levels in our study area occur mainly within natural populations of *E. cazorlanum* and *E. cicutarium,* with stronger population structure in *E. cicutarium,* and highlight the relevance of analyzing short distance spatial patterns.

## 1. Introduction

Understanding the sources of natural phenotypic variation is a crucial question in ecology and evolution. The idea that epigenetic diversity can compensate for reduced genetic diversity, promoting phenotypic variation through trait plasticity in response to environmental changes is relatively new (Balao et al., 2018; Bossdorf et al. 2008; Richards et al., 2017; Richards, 2006; Williams et al., 2023). Trait plasticity can be particularly advantageous for sessile organisms such as plants, with limited escape strategies to face environmental perturbations and, thus, the magnitude and population structure of genetic and epigenetic diversities in plant species with disparate genomic and ecological features warrants comprehensive investigation (Anderson & Gezon, 2015; Herrera et al.,2022; Richards et al., 2017; Szukala et al., 2023). Such knowledge can help to optimize decisions in conservation (i.e. accurately evaluating the adaptive potential of populations, Lamka et al., 2022; Williams et al., 2023), or even predict how plant species would fare when introduced in foreign ecosystems (i.e., assess invasive potential, Hawes et al., 2018).

DNA methylation remains a focus of interest among the spectrum of plant epigenetic modifications. The mechanistic control and effects of DNA methylation, its relationship with other chromatin regulators such as histone tail modification and small RNAs, and the potential transgenerational transmission of different methylation-marker types have been mainly studied in model species such as *Arabidopsis thaliana* (Cokus et al., 2008; Muyle et al., 2022; Stroud et al., 2014), and economically important species such as maize and rice (reviewed by Varotto et al., 2022; but see also Herrera et al., 2018). Several DNA-methyltransferases set methyl groups in the carbon-5’ of cytosines followed by specific nucleotide motifs in the genome, referred to as methylation contexts. In plants, there are three cytosine methylation contexts: CpG, CHG and CHH (where H can be any nucleotide but G), which are each regulated by the action of distinct enzyme complexes, and appear unevenly represented in plant genomes (Law & Jacobsen, 2010). Symmetrical methylation in CpGs can be preferentially found in promoters and exons within genes (referred to as gene body methylation; Bewick & Schmitz, 2017) and can be more frequently heritable compared to other contexts (Schmitz et al., 2011). In turn, methylated CHGs and CHHs are more abundant in heterochromatic and intergenic regions, and are often reset between generations or rapidly (de)methylated in response to changes in environmental conditions (Bouyer et al., 2017; Du et al., 2012). Furthermore, transposable elements (TEs) show DNA methylation in all three contexts, resulting in TE silencing and promoting genome stability (Matzke & Mosher, 2014). Given the wide range of genome sizes in plants, and the positive relationship between genome size and TE proportion in the genome (Kidwell, 2002), plant species show a broad variation in the level of DNA methylation, that was found to be related to phylogenetic relationships and species genome size (Alonsoet al., 2019a; Alonso et al., 2015; Vidalis et al., 2016).

The genomic and ecological features that could be associated to the broad variation in levels of DNA methylation mentioned above are still poorly understood. Particularly for non-model plant species, contrasting evidences have been found regarding their variation across natural populations and geographic-climatic gradients (see e. g. De Kort et al., 2020; Galanti et al., 2022; Lamka et al., 2022; Richards et al., 2017; Richards et al., 2010). In particular, global DNA methylation tends to be lower in woody plants than perennial herbs, as well as in genomes of widespread species compared to species with restricted geographic distributions (Alonso et al., 2019a). Relevance of such life history traits when controlling for phylogenetic relatedness remains largely unexplored and, thus, studies focused on analyzing natural variation in DNA methylation of related species with contrasting ecological features should be particularly valuable to better understand epigenomic similarities and divergences within and across plant species (Medrano et al., 2020 and references therein; Noshay & Springer, 2021). Reduced representation bisulfite DNA sequencing (RRBS) allows characterization of DNA methylation from a reduced portion of the genome and can be applied to species even when no reference genome is available, as it is usually the case for plants with narrow distribution ranges (Paun et al., 2019). Despite limitations, RRBS can provide quantitative methylation information at the different sequence contexts (CpG, CHG, CHH), together with an estimate of methylation similarity across individuals and differentiation between populations and species, and may lead to some functional characterization of methylated regions when an annotated reference genome of close relatives and/or complementary RNAseq data is available (Chen et al., 2015; Hsu et al., 2017; Kreutz et al., 2020; Nunn et al., 2021; Paun et al., 2019).

In this study, we explore the DNA methylation variation in natural plant populations of two *Erodium* species co-occurring in Cazorla mountain ranges (South East Spain): the endemic woody perennial *E. cazorlanum*, and the widespread annual herb *E. cicutarium*. Given the differences in habitat specialization, distribution range and life history of these two congeneric species, we predict they could differ in their genome-wide DNA methylation level and in the way epigenetic variation is structured within and among populations throughout a geographical area where both species co-exist. In particular, we predict that (1) individuals of the endemic *E. cazorlanum* will exhibit higher diversity in genomic methylation levels than genetic variants, and (2) that epigenetic variability should be less structured in space in the endemic species compared to its widespread congener given the expected reduced range in suitable environmental conditions for the former. To address so, we generated Restriction site-Associated DNA sequencing (RADseq) data, both untreated and treated with bisulfite (bsRADseq, Trucchi et al., 2016), from leaf DNA of the two *Erodium* species, each collected from three different localities. First, we checked for population genetic structure in each species using the single nucleotide polymorphisms (SNPs) obtained from RADseq data. We then estimated the percentage of DNA methylation per cytosine in the three methylation contexts to search for quantitative and qualitative differences in genome-wide DNA methylation at the species and locality levels using the bsRADseq data. Next, we identified differentially methylated cytosines (DMCs), used them to estimate population epigenetic structure in each species, and summarized them in differentially methylated regions (DMRs) to characterize genomic loci potentially under divergent epigenetic regulation, at locality and species level. Challenges derived from using a single incomplete reference genome to analyze both species are discussed at the end.

## 2. Materials and methods

### 2.1. Study species and sample collection

*Erodium cazorlanum* (Heywood) is an endemic, long-lived woody perennial plant, typically associated to a few dolomite outcrops in Sierra de Cazorla-Segura-Las Villas Natural Park (Jaén, SE Spain) (Alonso & García-Sevilla, 2013). *Erodium cicutarium* (L.) L’Hér. is a widespread, herbaceous annual plant species with a fast life-cycle, frequently associated with ruderal habitats, and to more developed soils within the same study area (Fiz-Palacios *et al.,* 2010). The genome size of *E. cicutarium* has been estimated to range from 1.2 to 2.7 Gbps (Pustahija et al., 2013; Zonneveld, 2019), different ploidy levels have been reported from different geographic regions, ranging from diploid to tetraploid (Pustahija et al., 2013; Zonneveld, 2019), and preliminary data suggest our plants have 2n = 40 chromosomes. Genomic information about *E. cazorlanum* is scarce, but its estimated genome size is 8.1 Gbps (Alonso *et al*. unpublished data), it is described as an octoploid (Alonso & García-Sevilla, 2013; Blanca et al., 2009) and preliminary data suggest our plants have 2n = 80 chromosomes.

Young fully-developed leaves from 18 *E. cazorlanum* and 18 *E. cicutarium* individuals were collected at flowering within three localities per species in Sierra de Cazorla-Segura-Las Villas Natural Park (Jaén, Spain, **Figure 1**). The number of individuals per locality was selected based on available information on minimum sample sizes required for accurate estimates of intra- and inter-population genetic diversity based on high-throughput DNA sequencing (Nazareno et al., 2017). The study localities (hereafter populations) were Fuente Bermejo (FB), Puerto Lézar (PL), and Puerto del Tejo (PT) for *E. cazorlanum*; and Nava Correhuelas (CH), Fuente Bermejo (FB), and Puerto del Tejo (PT) for *E. cicutarium*. Horizontal distances between the nearest and farthest sampling sites were 7.5 and 19.4 km for *E. cazorlanum* (FB-PL and PL-PT, respectively), and 2.1 and 11.5 km for *E. cicutarium* (CH-FB and FB-PT, respectively). Note that despite sharing some sampling localities, the two species were located in separated patches due to differences in soil requirements (Moreno, 2008). Sampling details are thoroughly described in (Medrano et al., 2020).

**Figure 1.**
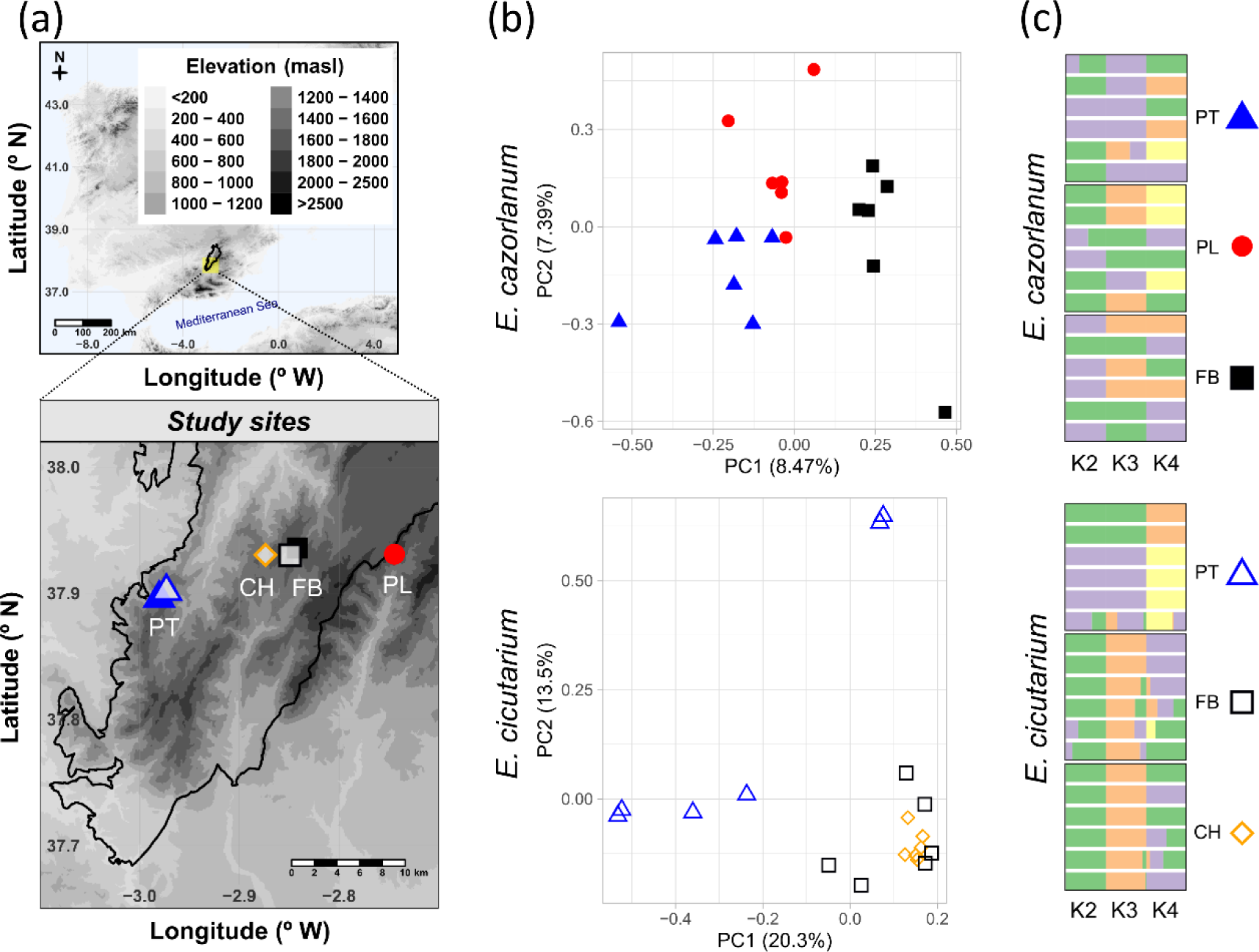
Genetic population structure of the two study species. A: Study area and localities. Top: map of the Iberian Peninsula showing the area of the Parque Natural Sierras de Cazorla-Segura-Las Villas (delimited by a solid black line) and the sampling area (yellow square). Altitude (masl = meters above sea level) is represented as a grey scale. Bottom: Detail of the sampling area, showing the study localities of *Erodium cazorlanum* and *E. cicutarium*. CH = Nava Correhuelas, FB = Fuente Bermejo, PL = Puerto Lézar, PT= Puerto del Tejo. B: Principal component analysis for *E. cazorlanum* (top) and *E. cicutarium* (bottom) based on SNPs obtained by RADseq. Each biplot shows principal components PC1 and PC2 (together with their percentage of explained variance) in X- and Y-axis, respectively; data points are individuals; populations are shown with different colors and shapes. C: Admixture analysis for *E. cazorlanum* (top) and *E. cicutarium* (bottom) based on SNPs from RADseq. X-axis shows the K value chosen for each model. Y-axis shows all study individuals, with populations enclosed in different boxes. Colors represent different modeled genetic lineages according to the selected K value. Each color within a tile indicates the proportion of SNPs from different modeled lineages.

### 2.2. DNA extraction, library preparation, sequencing and quality check

Total genomic DNA was extracted using Bioline ISOLATE II Plant DNA Kit and quantified using a Qubit fluorometer 2.0 (Thermo Fisher Scientific, Waltham, MA, USA). The RADseq library preparation followed the general steps from (Balao et al., 2023) (see **Supplementary Materials & Methods** for details). Half the DNA yield per sample was treated with bisulfite to generate bsRADseq libraries (Trucchi et al., 2016), the other half remained untreated for generating RADseq libraries.

Both untreated and bisulfite-treated DNA samples were sequenced at the VBCF Vienna (https://www.viennabiocenter.org/vbcf/next-generation-sequencing/) as paired-end 2 x 150 bp reads, a single lane of an Illumina HiSeq 2500 machine was used for the untreated samples of the two species (RADseq) and another lane was used for the bisulfite-treated samples (bsRADseq). Reads are accessible through the sequence read archive (SRA) from NCBI, under BioProject ID PRJNA1071520. The reads were demultiplexed with deML (Renaud et al., 2015), and then STACKS v2.41 (Rochette & Catchen, 2017), discarding reads with low quality scores according to default parameters. Demultiplexed reads were trimmed and cleansed from Illumina adapter sequences with Trimmomatic v0.39 (Bolger et al., 2014) using the options ‘*TrimmomaticPE-phred33 ILLUMINACLIP:adapterList:2:30:10 LEADING:28 TRAILING:28 SLIDINGWINDOW:4:15 MINLEN:30*’.

### 2.3. Population genetics analysis

A draft genome of *E. cicutarium* based on PacBio single-molecule long-read sequencing and Illumina short-reads sequencing was assembled and annotated (see **Supplementary Materials & Methods** for the details). The draft genome v2 is available at NCBI Genome, under accession JBDFRG000000000; PacBio reads are available at NCBI SRA, under BioProject ID PRJNA984161. Trimmed reads from the RADseq libraries were aligned to the *E. cicutarium* draft genome v2 using *bowtie2* v2.2.5 (Langmead & Salzberg, 2012) with the ‘*--very-sensitive*’ option. Mapped reads were analyzed with the STACKS script *ref_map.pl*, and then single nucleotide polymorphisms (SNPs) were filtered using STACKS binary *populations* with options ‘*-r 0.83 --min_maf 0.05 -- max_obs_het 0.7 --vcf --write_single_snp*’. For each species separately, remaining SNPs were analyzed with PLINK v1.90b7 (Purcell et al., 2007) to perform Principal Component Analysis (PCA) to summarize the genomic variation. Further, each SNP dataset was analyzed with ADMIXTURE v1.3.0 (Alexander & Lange, 2011) to test the population genetic structure of each species, testing for K values from 1 to 6, choosing the best fit for the K value with the lowest cross-validation error. Pairwise genetic distance matrices between individuals within a species were generated from the SNP dataset using the *distance* function of PLINK, that calculates pairwise identity-by-state (IBS) distances based on allele counts from alleles shared for each pairwise comparison. As an alternative approach, additional analyses for SNP calling and population structure were performed with different model assumptions as implemented in the genotype likelihood-based methods of *ANGSD* (Korneliussen et al., 2014) and *NGSadmix* (Skotte et al., 2013) (see **Supplementary Materials & Methods** for details). Despite being developed for diploid organisms, ANGSD was suggested as a powerful approach to disentangle genetic patterns in heteroploid datasets (Zaveska et al., 2019). Naturally occurring C/T polymorphic sites found in any of the two RADseq dataset were masked in the *E. cicutarium* genome for downstream analyses.

### 2.4. Alignment of bisulfite converted reads to genome

Trimmed reads from the bsRADseq were aligned to the same *E. cicutarium* draft genome using *bismark* v0.23.1 (Krueger & Andrews, 2011), with *Bowtie2* as base aligner (Langmead & Salzberg, 2012), using the following options: *‘--bowtie2 --non_directional --most_valid_alignments 2 --L 32 --D 10 --R 1 --score_min L,0,0.2’*. For each individual, we checked the summary report recording the mapping efficiency, the number of cytosines screened, and the distribution among the different contexts (CpG, CHG, and CHH). The next step was to extract the methylation information of each cytosine position using *bismark_methylation_extractor*, with the options *‘--no_overlap --ignore 4 –report --bedGraph --counts --CX’*. As in each read pair, the read corresponding to the sonicated end incorporates an unmethylated cytosine in its first position during library construction, which then is exposed to bisulfite treatment, bisulfite conversion rate (BCR) was calculated for each sample using the formula *100 * T / (T + C)*, where *T* and *C* are the respective percentage of thymines and cytosines in the first position of the 2^nd^ paired read (Krueger, 2013).

### 2.5. Global DNA methylation analysis

Unless otherwise stated, all statistical analyses were performed using R v4.1.2 (R Core Team, 2023). The number of methylated cytosines and total number of genomic positions analyzed per individual was extracted from *bismark* reports with a custom R script. Global effects associated to species and population divergence on global genome methylation levels were analyzed with a GLM with quasibinomial distribution and logit link, using *emmeans* (Lenth, 2023) and *lme4* (Bates et al., 2015) R packages. The dependent variable was the overall proportion of methylated to unmethylated cytosines per individual, and the formula included species as fixed factor and populations of each separate species as a nested fixed factor (formula: proportion of methylated cytosines ∼ species + species: population). The same process was done for each methylation context separately (CpG, CHG, CHH).

### 2.6. Differential methylation analysis

Detection of differentially methylated cytosines (DMCs) between populations was conducted with *DSS* v2.46.0 R package (Feng et al., 2014; Park & Wu, 2016). *DSS* is based on a beta-binomial regression model with arcsine link function, and it takes advantage of shrinkage estimation of the dispersion parameter based on a Bayesian hierarchical model to reduce the dependence of variance on mean. P-values were adjusted for multiple testing using the Benjamini and Hochberg false discovery rate (FDR) procedure, retaining those with FDR < 0.05 (Benjamini & Hochberg, 1995). Analyses were conducted for each species separately. Although *DSS* allowed fitting a model with a single factor with three populations as levels, handling RRBS data reduces the probability of finding single cytosine positions well covered for all three populations (i.e., a position showing acceptable coverage for two populations, but not covered in the remaining population), which would discard potential DMCs due to insufficient coverage. Thus, differences in cytosine methylation between pairs of populations were calculated fitting the data to one-factor models consisting in pairs of populations, generating six datasets (FB x PL, FB x PT and PL x PT for *E. cazorlanum*; CH x FB, CH x PT and FB x PT for *E. cicutarium*). A position was considered a DMC when its FDR between populations was < 0.05, and the difference in methylation was > 20 % between any of the paired-population comparisons. This approach implies that the definition of hyper- and hypo-methylation is relative: a single DMC can be considered as hyper-methylated in one or more populations, and at the same time hypo-methylated in other population(s). Number of overlapping DMCs (either hyper- or hypo-methylated DMCs) from two populations were analyzed with a hypergeometric test to investigate whether overlaps were greater than expected by chance.

Hierarchical clustering and visualization based on methylation percentage per DMC was calculated and plotted using *pheatmap* R package (Kolde 2019). The variation observed in percentage of methylation across DMCs was also analyzed with non-metric multidimensional scaling (NMDS) in *vegan* R package (Oksanen et al., 2022), with 100 random starting configurations and choosing the Manhattan distance method. Sets of DMCs which showed 100 % methylation in a single individual and methylation percentages close to zero in the rest of the individuals were named “drastic” DMCs, the rest of DMCs were named “gradual” DMCs. Hierarchical clustering and NMDS analyses were performed again with only “gradual” DMCs.

Epigenetic Euclidean distance matrices between individuals within a species were calculated from the methylation percentage matrix of gradual DMCs using *vegan* R package (Oksanen et al., 2022). The complete DMC set and only “gradual” DMC set epigenetic matrices were compared with previously calculated genetic distance matrices with a Mantel test, to test for intra-specific Spearman correlation between genetic and epigenetic distance, with 9,999 permutations, using *vegan* R package. Additional Mantel’s tests to check the relationship between genetic and epigenetic distances were also performed on each separated methylation context (CpG, CHG and CHH).

Differentially methylated regions (DMRs) between populations were detected using *callDMRs()* function from the *DSS* package with default parameters. Methylation site density (MSD) per DMR was calculated by dividing the number of DMCs found in the DMR by the total length in base pairs of the respective DMR. After observing the MSD frequency distribution, percentile 90 of each population comparison’s MSD distribution was calculated and used as threshold to filter potential false positive DMRs (i. e. long DMRs with a relatively small number of DMCs). A DMR was retained when three or more DMCs were present within a genomic region with a minimum length of 50 bps and when its MSD value was higher than the percentile 90 of the MSD distribution for the analyzed comparison. After considering the biological differences of CpG, CHG and CHH methylation contexts, we opted to call DMRs as groups of DMCs from the same methylation context (CpG, CHG and CHH).

### 2.7. Inter-species comparison

The mean percentage of uniquely mapped bsRADseq reads to *E. cicutarium* draft reference genome was 19.00 % [13.6 % - 25.6 %] for *E. cazorlanum* samples, and 58.40 % [47.2 % - 72.6 %] for *E. cicutarium* samples (**Supplementary Table S1**), which showed statistically significant differences (ANOVA F statistic = 515.91, df = 1, P-value < 0.001). Given the different mapping rates, biases in differential methylation analysis were expected when cytosine and methylated cytosine coverages were compared between species. Thus, a normalization factor was applied to *E. cicutarium* read counts before comparing methylation levels between species. The normalization factor was calculated as the mean mapping efficiency from *E. cazorlanum* samples divided by the mean mapping efficiency for *E. cicutarium*, and *E. cicutarium* read counts were normalized by multiplying both the coverage and methylated cytosines counts per locus by the obtained normalization factor. Raw read counts in *E. cazorlanum* and normalized read counts in *E. cicutarium* were used for DMC and DMR calling analyses in the interspecies comparison. Priors for considering a DMC and a DMR were the same as the ones explained in previous sections above.

### 2.8. Functional characterization of differentially methylated regions

DMCs included in DMRs that overlapped either with exons, introns, 5’ UTRs, 3’ UTRs or intergenic regions as described in the *E. cicutarium* draft genome were identified with custom scripts. DMCs that overlapped with repeated elements from the *E. cicutarium* genome, including transposable elements (TEs), were retrieved with the *CNEr* R package (Tan et al., 2019). Fisher’s exact test was used to test whether the proportion of TEs overlapping with the *E. cicutarium* set of DMCs/DMRs was over- or under-represented compared to the proportion of all the TEs present in the *E. cicutarium* draft genome.

GO terms were extracted from the gene annotation file of the *E. cicutarium* draft genome with a custom script. Lists of DMRs overlapping with annotated genes were then analyzed with *topGO* (Alexa & Rahnenfuhrer, 2023) to find enriched GO terms in each DMR list, using a weighted Fisher’s exact test and correcting the p-value with FDR. A GO term was considered enriched/over-represented in a set when the FDR < 0.05.

## 3. Results

### 3.1. Sequencing quality and bisulfite treatment efficiency

For the 36 samples, Illumina sequencing produced a total of 332.2 million 2 x 150 bp length paired-end reads for RADseq (ranging from 3.4 to 12.2 million per sample in *E. cazorlanum*, and from 4.4 to 33.8 million per sample in *E. cicutarium*), and a total of 323.4 million 2 x 150 bp length paired-end reads for bsRADseq (ranging from 3 to 13.6 million per sample in *E. cazorlanum*, and from 3.8 to 30 million per sample in *E. cicutarium*). After removing adapters and checking the distribution of sequencing quality per nucleotide, 315.3 million reads were retained in the RADseq, and 302.4 million reads remained in the bsRADseq. Mean bisulfite conversion rate per sample was 99.09 % [99.07 % - 99.13 %] (**Supplementary Table S1**).

### 3.2. Genetic polymorphism and population structure

A total of 24,977 SNPs in *E. cazorlanum* and 31,696 SNPs in *E. cicutarium* passed the filtering process. In the endemic *E. cazorlanum*, the PCA shows clustering by sampling locality mainly on PC1, but the position of the groups did not follow a geographical pattern, with some individuals from different localities positioned very close together in the multidimensional PCA. Individuals from the FB population appeared more clearly separated from the other two populations, although all principal components explained less than 10 % of the variance on their own (**Figure 1**). ADMIXTURE considered *E. cazorlanum* as a single genetic entity (K = 1) based on minimizing the cross-validation error, whereas higher numbers of potential populations did not differentiate between the different localities. For the widespread *E. cicutarium*, the PCA shows three separated groups: one with all the individuals from CH and FB populations, another with four individuals from PT population, and the third one with two individuals from PT population (**Figure 1**); the difference between the first two groups is supported by PC1 (20.3 % of explained variance) and the difference between the first and third group is described by PC2 (13.5 % of explained variance). ADMIXTURE again considered *E. cicutarium* individuals as a single genetic entity (K = 1) based on the cross validation error (**Supplementary Figure S1**). **Figure 1** shows assignment of *E. cazorlanum* and *E. cicutarium* individuals to subpopulations, assuming models with two to six subpopulations. **Supplementary Table S2** shows the cross-validation error values. Alternative analyses performed with the genotype likelihood-based method of *ANGSD* showed consistent results with the ones shown above (see **Supplementary Materials & Methods** for details).

### 3.3. Global DNA methylation analysis

*Bismark* analyzed a total of 11,921,613 cytosines in any methylation context (1.49 % of *E. cicutarium* draft genome total length). Percentages of global DNA methylation ranged from 6.66 % to 12.71 % in the endemic *E. cazorlanum* and from 7.76 % to 11.07 % in the widespread *E. cicutarium* (**Supplementary Table S1**). Within species variance of global methylation percentages was higher for *E. cazorlanum* (**Table 1**). **Figure 2** shows the methylated cytosine percentages per sequence context by species and by population. On average, there were significant differences between the two species in the ratio of methylated to unmethylated cytosines per individual (**Table 2**) with the genomes of *E. cazorlanum* being more methylated (mean ± SD = 9.51 ± 1.83) compared to *E. cicutarium* (mean ± SD = 9.08 ± 0.94). Similar results were also obtained for each of the three methylation contexts (**Table 2**, **Supplementary Figure S2**). Differences in percentages of global DNA methylation were also statistically significant between the study populations within a species, as were all pairwise within-species population contrasts (**Table 2**, **Supplementary Figure S2**). Further, within species variance was always higher for *E. cazorlanum* global and context specific methylation percentages (**Table 1**).

**Figure 2.**
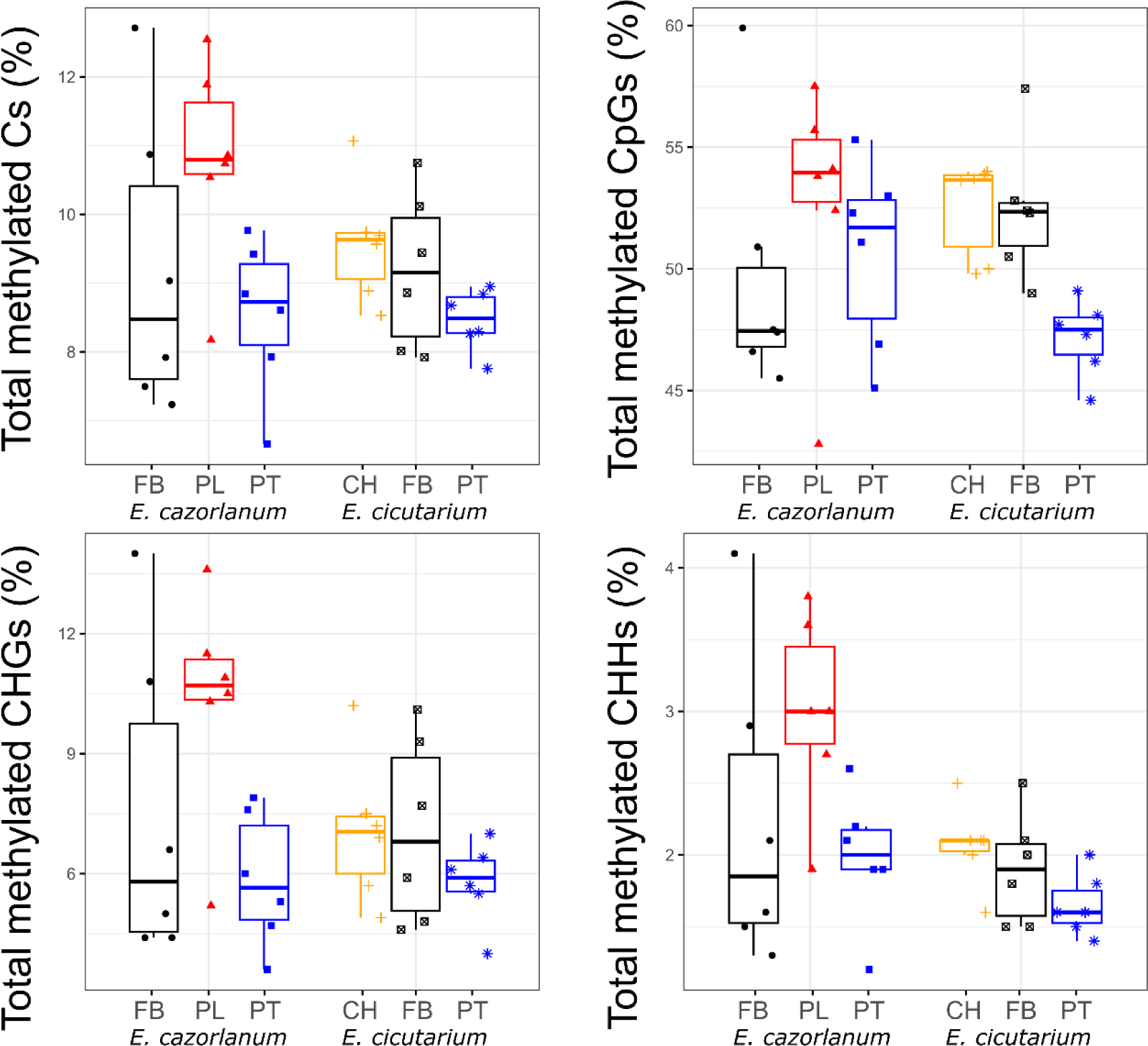
Variation in levels of DNA methylation in populations of *E. cazorlanum* and *E. cicutarium*. The plots show percent of all the sequenced cytosines that were methylated (top left) and estimates obtained for cytosines in the three methylation contexts: CpG (top right), CHG (bottom left) and CHH (bottom right). On each graph, symbols represent individuals, X-axis shows the study populations of the two *Erodium* species, and Y-axis shows the methylation percentage estimated. Each box shows the median (central line), quartiles 1 and 3 (bottom and top edges of the box), and maximum and minimum values (box whiskers) per population (represented by different colors and shapes). Note that Y-axes have different scales.

**Table 1.** Coefficients of variation of global DNA methylation percentages. First column shows each methylation context. Each set of four columns shows information related to each species. The first column of each species set shows CV from all populations, the three remaining columns show CVs for each population.

| CVs | *E. cazorlanum* |  |  |  | *E. cicutarium* |  |  |  |
| --- | --- | --- | --- | --- | --- | --- | --- | --- |
|  | Overall | FB | PL | PT | Overall | CH | FB | PT |
| Total Cs | 18.72<br>% | 22.54<br>% | 13.59<br>% | 12.54<br>% | 9.74% | 7.55% | 12.30<br>% | 5.41% |
| CpG | 9.28% | 10.77<br>% | 9.79% | 7.65% | 6.51% | 3.85% | 5.42% | 3.34% |
| CHG | 42.69<br>% | 52.90<br>% | 26.94<br>% | 28.58<br>% | 27.24<br>% | 25.79<br>% | 32.99<br>% | 17.68<br>% |
| CHH | 35.50<br>% | 47.68<br>% | 22.61<br>% | 23.31<br>% | 17.93<br>% | 13.91<br>% | 20.25<br>% | 13.14<br>% |

**Table 2.** Global cytosine methylation differences at species and population level. First column shows the set of cytosines for which a binomial distribution model was run to test whether the difference between the proportion of methylated to unmethylated cytosines (mC / uC) was statistically significant. Columns two to four show log-ratio χ2, degrees of freedom, and p-value for species. Columns five to seven show the same for populations within species.

| Proportion<br>of mC / uC | Species |  |  | Species:Population |  |  |
| --- | --- | --- | --- | --- | --- | --- |
| | LR $\chi^2$ | d.f. | P-value | LR $\chi^2$ | d.f. | P-value |
| Total Cs | 74,280 | 1 | < 0.001 | 918,941 | 4 | < 0.001 |
| CpG | 2,118 | 1 | < 0.001 | 627,781 | 4 | < 0.001 |
| CHG | 146,988 | 1 | < 0.001 | 498,203 | 4 | < 0.001 |
| CHH | 340,129 | 1 | < 0.001 | 426,370 | 4 | < 0.001 |

### 3.4. Differential cytosine methylation analysis within species

A total of 12,795 and 16,131 naturally occurring C/T polymorphic sites obtained from RADseq analyses were masked in bsRADseq data before analyses in *E. cazorlanum* (0.11 % of analyzed Cs) and *E. cicutarium* (0.14 % of analyzed Cs), respectively. For *E. cazorlanum*, a total of 13,643 DMCs (5,632 CpG, 2,605 CHG, 5,406 CHH) were detected between the three populations (ca. 0.11 % of total analyzed Cs), and 47,615 DMCs (ca. 0.40 % of total analyzed Cs) were detected between the three populations of *E. cicutarium* (20,670 CpG, 8,278 CHG, 18,667 CHH; **Supplementary Figure S3**, **Supplementary Tables S3, S4, S5**). In each of the two species, the number of overlapping hyper- or hypo-methylated DMCs between each population pair was never significantly greater than expected by chance (hypergeometric test, P-value = 1, **Supplementary Figures S4, S5**). The overlap of DMCs between species was also not greater than predicted by chance either (hypergeometric test, P-value = 1, **Supplementary Figure S6**).

Hierarchical clustering of DMCs showed that most clusters were defined by cytosines with “drastic” changes in methylation percentage within the study dataset (i. e., 100 % methylation for one individual and ca. 0 % methylation for the rest, **Supplementary Figure S7**). Removing these “drastic” DMCs and leaving only those with “gradual” change in methylation percentage still did not group samples following a clear population structure in any of the two species (**Figure 3, Supplementary Figure S7**). When analyzing the “gradual” DMCs by separated methylation context, the CpG context DMCs showed clusters with at least three samples from the same population for *E. cazorlanum*, and more patently for *E. cicutarium*; while CHG and CHH contexts did not group more than three samples from the same population within any cluster (**Supplementary Figures S8, S9**).

**Figure 3.**
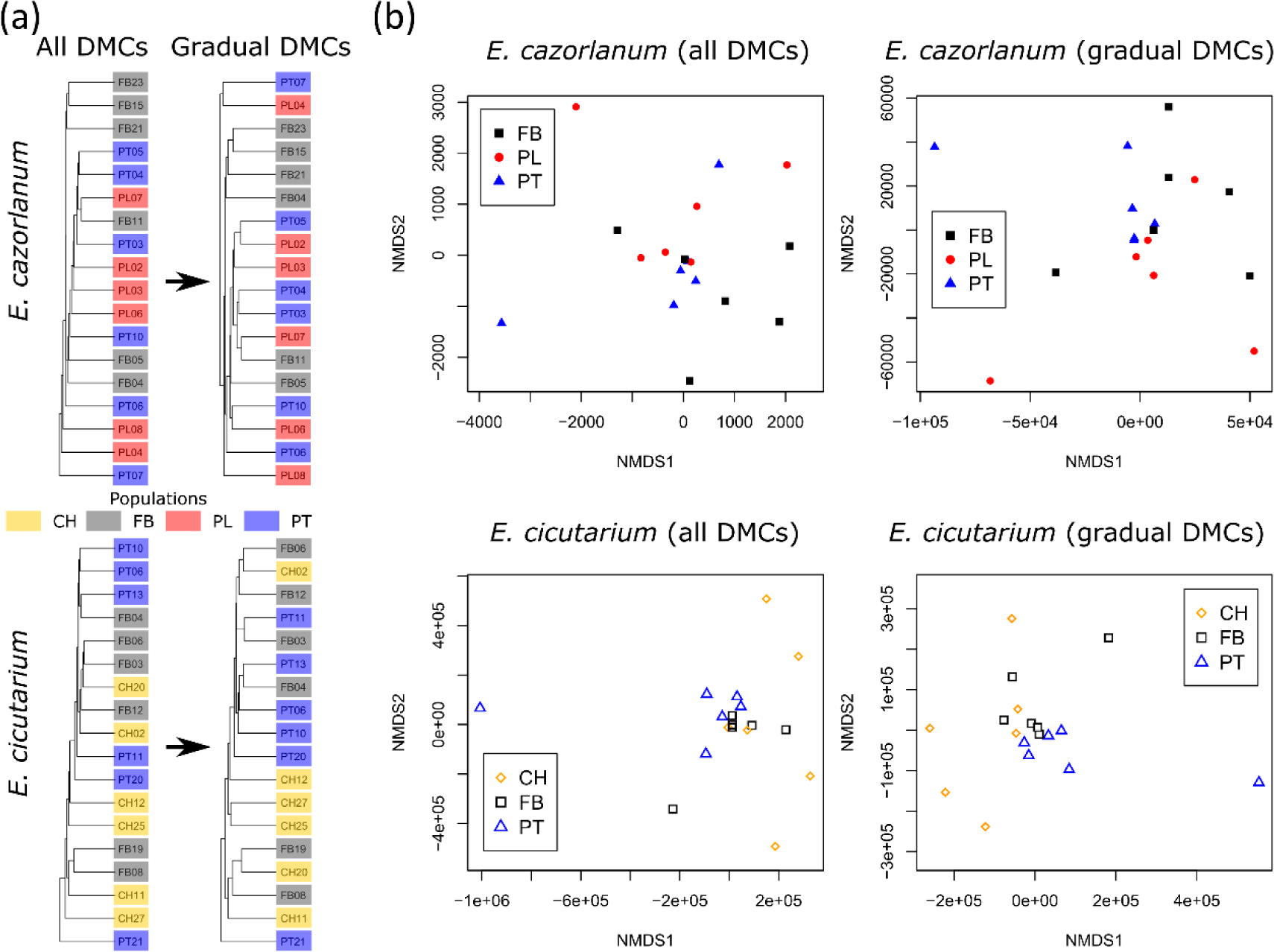
Epigenetic population structure of the two *Erodium* species. A: Hierarchical individual clustering according to obtained differentially methylated cytosines (DMCs) based on percentage of methylation per DMC locus. Top dendrograms correspond to *E. cazorlanum* (N = 13,643 loci) and bottom dendrograms to *E. cicutarium* (N = 47,615 loci). Each column shows the resulting clustering of all DMCs (left) and gradual DMCs (right), see text for details. Colored tiles attached to each sample ID show the population of origin. B: Multilocus analysis of individual variation in methylation percentage of all DMCs (left graphs) and gradual DMCs (right graphs) based on non-metric multidimensional scaling (NMDS). Top graphs show data for *E. cazorlanum*, bottom graphs for *E. cicutarium*. X and Y axes show non-linear multidimensional scaling (NMDS) dimensions NMDS1 and NMDS2, respectively. Colors of data points show the population of origin.

Multivariate analysis based on all DMCs showed no clear population segregation in *E. cazorlanum* and a slight differentiation of the PT population for *E. cicutarium* (**Figure 3**). The same analysis, but performed only with gradual DMCs, marginally increased the differentiation of PT population from FB and CH in *E. cicutarium*, while not affecting *E. cazorlanum* grouping pattern (**Figure 3**). Performing multivariate analyses with DMCs separated by context did not improve segregation of populations (**Supplementary Figures S10, S11**).

Mantel tests revealed no statistically significant correlation between pairwise genetic and epigenetic distances across individuals in any of the two study species (*E. cazorlanum*: Mantel’s R = 0.104, p-value = 0.271; *E. cicutarium*: Mantel’s R = 0.003, p-value = 0.471). When considering epigenetic distances exclusively for each separate methylation context, Mantel tests also showed no statistical significance (p-value > 0.05).

### 3.5. Genomic and functional characterization of DMRs

Only DMRs with high density of DMCs (defined as DMRs above the percentile 90 of the MSD distribution; **Supplementary Figure S12, Supplementary Table S6**) were selected for further characterization. **Supplementary Table S7** shows the full list of DMRs that passed the filter, including a total of 43 DMRs defined between populations of the endemic *E. cazorlanum* that vary in length between 53 and 165 bp, and 145 DMRs defined between populations of the widespread *E. cicutarium* that vary in length between 51 and 343 bp. GO term enrichment analysis did not reveal enriched GO terms in any set of DMRs. In *E. cazorlanum*, we found 19 DMRs whose available annotation was associated with genes involved in plant development, amino acid metabolism, or stress response. In *E. cicutarium*, 49 out of 145 DMRs showed available annotation, and were associated with plant development, stress response, signaling, and cell wall homeostasis, among other biological processes.

The two species showed an overall similar pattern regarding DMR overlapping with genomic features (**Figure 4**). DMRs in CpG context (17 in *E. cazorlanum*, 18 in *E. cicutarium*) overlapped most frequently with exons in both species (93.8 % in *E. cazorlanum*, 61.1 % in *E. cicutarium*), while DMRs in CHH context (24 in *E. cazorlanum*, 125 in *E. cicutarium*) overlapped evenly with exons, introns and intergenic regions. The number of DMRs in CHG context (three in *E. cazorlanum*, two in *E. cicutarium*, all overlapping with exons) is too low to reach any clear conclusion. Altogether, from the 43 DMRs found in *E. cazorlanum*, 29 (67.4 %) overlapped with exons, six (14.0 %) did so with introns, four (9.3 %) with intergenic regions, two overlapped simultaneously with an exon and an intron (4.7 %), one with a 5’UTR (2.3 %), and one (2.3 %) with a 3’ UTR. We found more evenly distributed overlaps in the 145 DMRs found between populations of *E. cicutarium*: 44 (30.6 %) overlapped with intergenic regions, 42 (29.2 %) with exons, 41 (28.5 %) with introns, eleven with an exon and an intron (7.6 %), four (2.8 %) with a 3’UTR, one with an intron and a 3’UTR (0.7 %) and one with a 5’UTR and an exon (0.7 %). A higher proportion of DMRs was found in intergenic regions in *E. cicutarium* (30.56 %) compared to *E. cazorlanum* (9.30 %).

**Figure 4.**
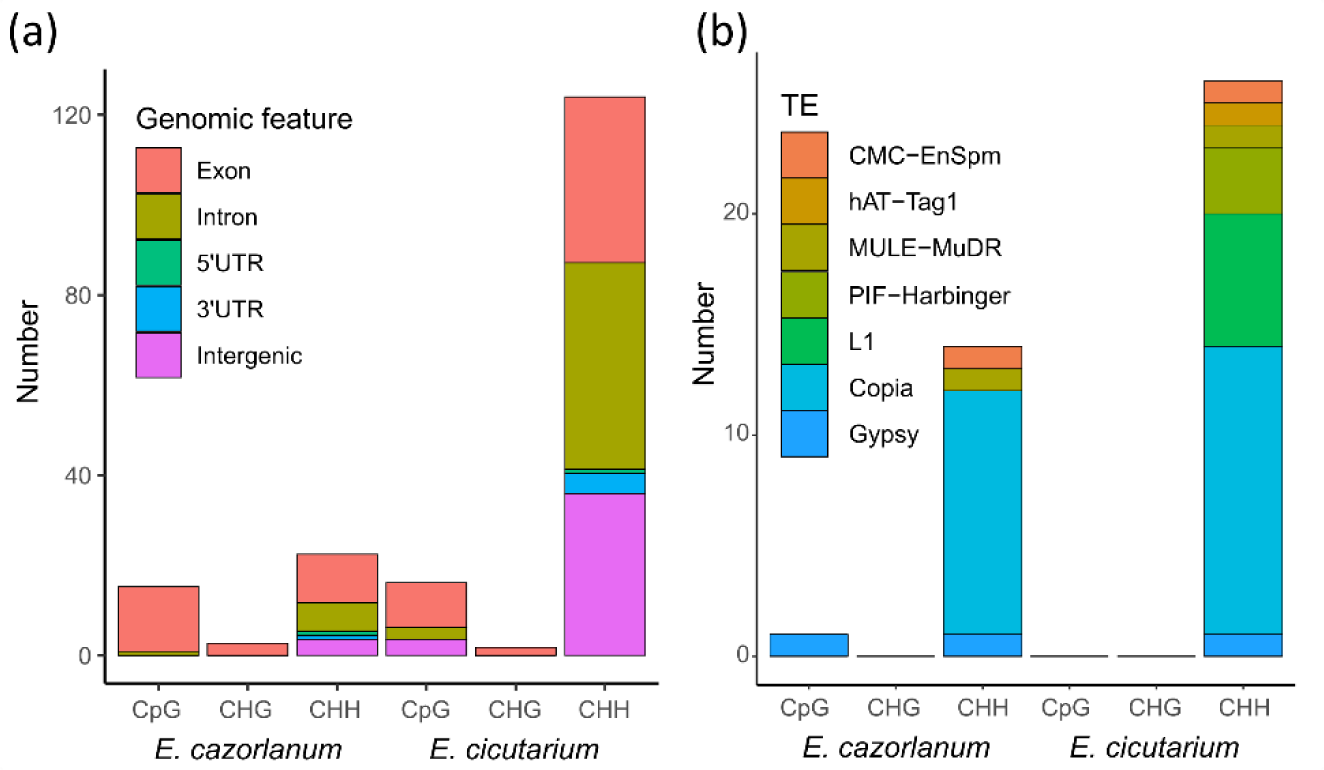
Genomic characterization of differentially methylated regions (DMRs) obtained for the two *Erodium* species. A: Genomic features overlapping with DMRs. X-axis shows the species and DMR methylation context (*E. cazorlanum*: CpG = 16, CHG = 3, CHH = 24; *E. cicutarium*: CpG = 18, CHG = 2, CHH = 125). Y-axis shows the numbers of DMRs found overlapping with each of the five different genomic features distinguished (exon, intron, 5’UTR, 3’UTR and intergenic regions). B: Transposable elements overlapping with DMRs. X-axis shows the species and DMR methylation context (*E. cazorlanum*: CpG = 1, CHG = 0, CHH = 14; *E. cicutarium*: CpG = 0, CHG = 0, CHH = 27). Y-axis shows the number of DMRs found overlapping with each of the nine families of TEs distinguished.

Regarding DMRs overlapping with transposable elements (TEs), most of them occurred in DMRs associated to CHH methylation context: 14 out of 15 DMRs overlapping with TEs involved methylation changes in CHH in *E. cazorlanum* (93.33 %), and all the 26 DMRs overlapping with TEs in *E. cicutarium* involved only methylation changes in CHH (**Figure 4**). In *E. cazorlanum*, TEs with most overlaps with DMRs were LTRs (with eleven *Copia* and two *Gypsy* elements overlapping with a DMR), followed by three DNA TEs (one *MULE-MuDR*, one *CMC-EnSpm*, and one uncharacterized DNA element overlapping with a DMR). For *E. cicutarium*, the most overlapped TE family was also *Copia* with 13 DMRs, in addition, it showed six *LINE/L1*, three *PIF-Harbingers*, one *MULE-MuDR*, one *hAT-Tag1*, one *CMC-EnSpm*, and one *Gypsy* element overlapping with DMRs. **Supplementary Table S8** shows all overlaps between a DMR with a genome feature, a TE, or both.

The set of DMRs from *E. cicutarium* showed a higher than expected proportion of overlaps with *Copia* TEs compared with the proportion of *Copia* TEs in the genome of *E. cicutarium* (χ^2^ = 3.522, df = 1, p-value = 0.045, **Supplementary Figure S13, Supplementary Table S9**). An analogous analysis in *E. cazorlanum* was not conducted because the TE set in *E. cazorlanum* genome may be substantially different from *E. cicutarium* genome, making the comparison between *E. cazorlanum* TEs in DMRs and *E. cicutarium* TE genome set inadequate.

### 3.6. Comparative analysis between species

Given the difference in read mapping efficiency between species (**Supplementary Figure 14**), we used the ratio between mean mapping efficiencies (0.33) as normalization factor when comparing *E. cicutarium* read coverage values with *E. cazorlanum*. First, we compared samples from *E. cazorlanum* and *E. cicutarium* before normalization of *E. cicutarium* read counts, detecting 10,765 DMCs (2,790 CpG, 2,272 CHG, 5,703 CHH) and 113 DMRs (12 CpG, 24 CHG, 77 CHH). After normalization, we found 13,744 DMCs (6,739 CpG, 2,154 CHG, 4,851 CHH, **Supplementary Table S10, Supplementary Figure S14**) and 183 DMRs (109 CpG, 23 CHG, 51 CHH), which were reduced to 19 DMR (5 CpG, 2 CHG, 12 CHH) after filtering by MSD (**Supplementary Table S11, Supplementary Figures S14, S15**). Similarity in the number of DMCs before and after normalization was significantly greater than expected for all contexts (hypergeometric test, P-value < 0.001), although the direction of the loss of DMCs was not the same for the three sequence contexts: 3,961, 232, and 351 DMCs were gained, while 12, 350 and 1,203 DMCs were lost after normalization in CpG, CHG and CHH contexts, respectively. Before and after normalization, *E. cicutarium* showed more hyper-methylated CpG sites compared to *E. cazorlanum*, while *E. cazorlanum* showed more hyper-methylation in CHG and CHH contexts compared to *E. cicutarium*.

Hierarchical clustering based on methylation percentage of the DMCs did not show a perfect divergence of species, but clades showed a species-like pattern rather than a population-like pattern, and the use of “gradual” DMCs did not change this pattern (**Supplementary Figure S16**). Multivariate analyses revealed a separation between centroids of the two species, a higher dispersion in *E. cicutarium* samples, and little within-species population variation, either using all DMCs or “gradual” DMCs (**Supplementary Figure S17**).

Among the 19 inter-specific DMRs obtained, we found association with an endochitinase (hydrolyses chitin and chitodextrins), *Cytohesin-2* (involved in regulation of protein sorting and membrane trafficking) and *pre-mRNA-splicing factor CEF1* (involved in cell cycle control) hyper-methylated in the CHH context in *E. cazorlanum*. The only DMR hyper-methylated in *E. cicutarium* was associated to the *LTL1* gene (involved in salt tolerance mechanisms) with increased methylation in the CpG context. Eleven DMRs from the inter-species comparison overlapped with exons, five with introns, two with an intergenic region, one with a 3’ UTR, and another one with both an exon and an intron. Ten DMRs from the inter-species comparison overlapped with fourteen TEs. They were mostly unknown repeats (eight out of 14), but also included three *Copia*, one *Gypsy*, one *SINE/tRNA-V*, and one *PIF-Harbinger* elements (**Supplementary Table S12**).

## 4. Discussion

This study analyzes variation in both DNA sequence (genetic) and cytosine methylation levels (epigenetic) at single-base resolution in natural populations of two *Erodium* species, the endemic *E. cazorlanum* and the widespread *E. cicutarium*, using bsRADseq and the structural and functional annotation provided by a draft reference genome developed *ad hoc* for the study species with the smallest genome (*E. cicutarium*). The two species exhibited sound differences in life history (woody perennial *vs*. annual), habitat specialization (dolomite specialist *vs*. ruderal) and geographic distribution range (local narrow endemic *vs*. world-wide widespread) that allowed us to test the hypothesis that population genetic and epigenetic structure within the region of coexistence would differ according to each species ecological features, with stronger epigenetic than genetic population structure in the least specialized (*E. cicutarium*). Besides, the use of methods that provide accurate genomic information of the methylation markers allowed the identification of a set of candidate genes whose epigenetic regulation might vary across the study localities of each species according to the recorded differences in DNA methylation, paving the avenue towards more specific analysis of adaptive strategies in species with contrasting genomic and ecological features. In the following paragraphs, we will first discuss the limitations of the reduced representation methodology used when applied to species with contrasting genome features and resources (see also Paun et al., 2019). Then, we will discuss the results obtained for our study system and how they reinforce the *a priori* prediction mentioned above.

### 4.1. Challenges and opportunities of RRBS analyses in plant ecological epigenetics

In absence of any high-quality gapless genome for any species in the *Erodium* genus, RRBS was the best available option to simultaneously detect natural genome and methylome variants across populations and explore their genomic location. In retrospective, six samples per population were estimated sufficient to reach an appropriate level of representation of what occurs in wild plant populations in genomics studies (Nazareno et al., 2017), and by using RRBS we expected to generate informative methylation datasets when conjointly used with genomic data for the same individuals (Gawehns et al., 2022; Paun et al., 2019). In fact, several studies have successfully identified differentially methylated loci by optimizing RRBS and its downstream analysis in commercial plant species with available reference genome (e.g., *Brassica rapa,* Chen et al., 2015; maize, Hsu et al., 2017). In this study, we succeeded in obtaining differentially methylated cytosines (DMC) and regions (DMR) between populations in the two *Erodium* species with six individuals per population, despite the limited power for differential methylation analysis presumed for the afforded sample sizes (Lea et al., 2017).

Interpretation of the results obtained for the portion of genome here explored should take into account that due to the relatively high GC content in the recognition site of the specific enzyme we used (*Pst*I that cuts at CTGCAG), bsRADseq dataset is enriched for genic regions (Lea et al., 2017; Trucchi et al., 2016) and, thus, quantitative global estimates would be less accurate for genomes with a relative high content of repetitive elements (i.e., in *E. cazorlanum* according to its larger genome size). Nevertheless, percentages of cytosine methylation per sequence context in the two species were consistent among each other and also with those reported for other plant species with complete methylomes, i.e., CpG higher than CHG, CHH being the lowest (Cokus et al., 2008; De Kort et al., 2020; Hsu et al., 2017; Zhao et al., 2020). The higher proportion of methylated to unmethylated cytosines obtained in *E. cazorlanum* is consistent with a marked difference in percentages obtained from independent analyses conducted by HPLC to obtain global DNA methylation estimates for the two species (28.4 % *vs*. 18.2 %; Alonso unpublished data). The sign of divergence in estimates of global DNA methylation per species could be associated to the largest genome size of *E. cazorlanum* (Alonso et al., 2015; Vidalis et al., 2016). The higher heterogeneity in overall methylation estimates of study individuals obtained for *E. cazorlanum* could be due to either genome size, and some unmeasured micro-environmental features (Valverde et al., 2024) or a combination of the two effects.

Further, using the reference genome from a close relative species generated a homology bias during the alignment of the reads that may weaken the potential for functional annotation of loci with divergent methylation in *E. cazorlanum*. Accordingly, the few DMRs obtained in *E. cazorlanum* were located in exonic regions, likely due to a higher similarity in the coding regions of the two species’ genomes and not necessarily to absence of methylation divergence in genomic regions that were not captured (Lea et al., 2017).

Altogether, our study exemplifies the challenges of applying NGS markers to species with limited genomic resources and complex genomes. However, conducting epigenomic studies of such species is compulsory to improve our current understanding of ecological factors behind the huge variation in global DNA methylation across plant species, that require a broad phylogenetic perspective (Alonso et al., 2015), and might be useful to better characterize the sources of variation in DNA methylation at plant ecosystem or community levels (Alonso et al., 2019b; Boquete et al., 2021). Our results support the use of RRBS for interspecific comparisons of natural variation in DNA methylation at difference sequence contexts, and a more restricted power to explore candidate genes and transposable elements potentially differentially regulated by DNA methylation when high-quality reference genomes are not available (Paun et al., 2019).

### 4.2. Genetic and methylation variation in natural populations of *E. cazorlanum* and *E. cicutarium*

In the small geographic area here explored, selected populations of each of the two study species lack or show limited signs of genetic population structure, suggesting they are well connected to each other, showing most genetic variation between individuals from the same population. Whereas this fact should be taken with caution for extrapolation to broader geographic areas, all study individuals could be associated to a single genetic background eventually, making the investigation of methylation variants more straightforward. Spontaneous epimutations, i.e. random changes in methylation status of cytosines, are estimated to be four to five orders of magnitude higher than mutation rate in *Arabidopsis*, which may facilitate the decoupling of methylation variation from genetic variation and some specific contribution to heritable phenotypic variation (Banta & Richards, 2018; Denkena et al., 2021; van der Graaf et al., 2015; Weng et al., 2019). In this study, we found absence of correlation between genetic and epigenetic distances across individuals in the two study species, supporting it would be worth to experimentally exploring their relative contribution to individual phenotypes and their plasticities under controlled environmental conditions (see e.g. Williams et al., 2023).

### 4.3. Genomic characterization of DNA methylation variants and functional annotation of DMRs

In each of the two species, most methylation changes between individuals with different population of origin were found in CpG and CHH contexts compared to a much lower frequency in CHG. A high level of CHH methylation variation is expected given the sensitivity of this context to environmental changes (Dubin et al., 2015); in fact, there is evidence that epigenetic distance based in CHH methylation differences can be positively correlated with climatic distance in *Thlaspi arvense* (Galanti et al., 2022). In the endemic *E. cazorlanum*, natural variation in methylation at those DMCs did not exhibit any population structure, regardless the analyses considered all or context-specific DMCs. In the widespread *E. cicutarium*, we found some epigenetic differentiation between the three study populations, supporting our expectation that, in absence of genetic population structure, the broader habitat conditions expected in a widespread species could contribute to larger methylation variance between study individuals at different scales (Valverde et al., 2024).

We only proceed with functional annotation of DMRs because solid experimental evidence has linked methylation gains and losses in large genomic regions (i.e., DMRs) with transcriptional variation and heritable phenotypic effects, whereas the functional relevance of single methylation changes (i.e., DMCs) might be more variable across plant species (De Kort et al., 2022; Denkena et al., 2021). Furthermore, although our analytical procedure based on *Bismark* concentrates exclusively on uniquely mapping reads that in turn will result in a depletion of representation of repetitive genomic elements, there was also a fair number of DMRs overlapping with TEs in our *E. cicutarium* data. As a rule, methylation levels in TEs are maintained high in plants to silence their expression and maintain genome stability (Matzke & Mosher, 2014). In our TE annotated bsRADseq data, the *Copia* family was the most abundant, and it was found overlapping with DMRs in a higher than expected proportion compared to the total number of TEs identified in the *E. cicutarium* draft genome. The expression of *Copia* TE family seems stress responsive in several plant species (Ito et al., 2011, 2016; Klein & Anderson, 2022), suggesting that *Copia* elements with CHH methylation islands in promoter regions that have shifted methylation between our study locations might be potentially linked to stress genes’ expression regulation. Additional studies in *E. cicutarium*, in which changes in DNA methylation caused by 5-azacytidine and the effect of recurrent drought were experimentally addressed, also found that induced DMRs were enriched with *Copia* elements (Balao et al., 2023), supporting the idea that this TE family may have a stress-related regulatory function in *E. cicutarium* too.

Finally, deep differences in read coverage obtained for the two species could cause biases in the direct inter-specific comparison of methylation landscapes offered by DMC detection. Following a normalization step to equilibrate coverage inspired in library size normalization for RNA-seq data (Conesa et al., 2016; Zeng & Mortazavi, 2012), we found that the overall number of detected DMCs between the two species was practically the same, although the relative frequency per sequence context changed and more significant methylation changes in CpG were retained after normalization. Further studies comparing two or more related species with a single available reference genome would be required to validate this methodology. Interestingly, main differences in genomic methylation profiles observed by DMC detection between the two study species involved hypermethylation in non-CpG sites in *E. cazorlanum* and hypomethylation of CpG sites, except when included in DMRs that were most frequently hypermethylated in all contexts.

### 4.4. Conclusions and future steps

With limited genomic resources typically available for non-model plants of conservation concern, this study compared DNA methylation at single-base resolution in a reduced portion of the genomes of two congeneric species and explored their spatial structure in a small area of coexistence. We found support for (i) an overall higher global DNA methylation in the endemic *E. cazorlanum*, particularly associated to hypermethylation in non-CpG sites, and in agreement with its higher genome size compared to its widespread congener *E. cicutarium*. (ii) The absence of genetic population structure in the endemic species, and (iii) the lack of correlation between genetic and epigenetic variation across individuals of both species, suggest that they could have independent effects on individual phenotypes. Finally, (iv) a slightly higher DNA methylation divergence between study populations in *E. cicutarium* supports the hypothesis that a stronger epigenetic, rather than genetic, population structure could be expected in widespread generalist species (see also Valverde et al. 2024). Future studies should take the advantage of the genomic resources generated here to link phenotypic, genetic and epigenetic variation in experimental settings suitable to identify the epigenetic contribution to plant adaptation to environmental variation.

## Author contributions

CA obtained funding for the study. MM, FB, OP, CA designed the experimental setting. MM and CA conducted fieldwork, PB performed molecular lab work. RM-B analyzed the data. RM-B wrote the first draft of the paper, with feedback from FB, OP and CA. All co-authors provided comments and suggestions on the final version of the manuscript and approved it.

## Acknowledgements

The authors gratefully thank to Noelia Zarza for assisting us during field sampling; to Ben Zonneveld (Netherlands) for determining the genomic sizes of *E. cazorlanum* and *E. cicutarium*; to Abelardo Aparicio (University of Sevilla, Spain) for determining the ploidy levels of *E. cazorlanum* and *E. cicutarium*; and to Carlos M. Herrera, Javier Valverde, Javier Puy and Anupoma N. Troyee (Estación Biológica de Doñana, Spain) for providing valuable discussion. Funding for this work has been provided by grant EPIENDEM (CGL2016-76605-P) from the Spanish Ministero de Economía y Competitividad, and grants EPINTER (PID2019-104365GB-I00) and DISTEPIC (PID2022-141530NB-C22) from the Spanish Ministerio de Ciencia e Innovación.

## Data accessibility and Benefit-generation statement

Illumina raw sequence reads are deposited in the SRA (BioProject PRJNA1071520). PacBio long reads are also deposited in the SRA (BioProject PRJNA984161). Draft genome assembly is deposited in NCBI Genome repository (accession JBDFRG000000000). Scripts are available under personal request to RM-B and CA. Benefits Generated: Benefits from this research accrue from the sharing of our data and results on public databases as described above.

## References

Alexa, B., & Rahnenfuhrer, J. (2023). topGO: Enrichment Analysis for Gene Ontology. Retrieved from https://bioconductor.org/packages/topGO doi:10.18129/B9.bioc.topGO

Alexander, D. H., & Lange, K. (2011). Enhancements to the ADMIXTURE algorithm for individual ancestry estimation. BMC Bioinformatics, 12, 246. doi: 10.1186/1471-2105-12-246

Alonso, C., & García-Sevilla, M. (2013). Strong inbreeding depression and individually variable mating system in the narrow endemic *Erodium cazorlanum* (Geraniaceae). Anales del Jardín Botánico de Madrid, 70(1), 72–80. doi:10.3989/ajbm

Alonso, C., Medrano, M., Perez, R., Canto, A., Parra-Tabla, V., & Herrera, C. M. (2019a). Interspecific variation across angiosperms in global DNA methylation: phylogeny, ecology and plant features in tropical and Mediterranean communities. New Phytologist, 224(2), 949–960. doi:10.1111/nph.16046

Alonso, C., Perez, R., Bazaga, P., & Herrera, C. M. (2015). Global DNA cytosine methylation as an evolving trait: phylogenetic signal and correlated evolution with genome size in angiosperms. Frontiers in Genetics, 6, 4. doi:10.3389/fgene.2015.00004

Alonso, C., Ramos-Cruz, D., & Becker, C. (2019b). The role of plant epigenetics in biotic interactions. New Phytologist, 221(2), 731–737. doi:10.1111/nph.15408

Anderson, J. T., & Gezon, Z. J. (2015). Plasticity in functional traits in the context of climate change: a case study of the subalpine forb *Boechera stricta* (Brassicaceae). Global Change Biology l, 21(4), 1689–1703. doi:10.1111/gcb.12770

Balao, F., Medrano, M., Bazaga, P., Paun, O., & Alonso, C. (2023). Long-term methylome changes after experimental seed demethylation and their interaction with recurrent water stress in Erodium cicutarium (Geraniaceae). bioRxiv. doi:10.1101/2023.01.19.524693

Balao, F., Paun, O., & Alonso, C. (2018). Uncovering the contribution of epigenetics to plant phenotypic variation in Mediterranean ecosystems. Plant Biology, 20 (S1), 38–49. doi:10.1111/plb.12594

Banta, J. A., & Richards, C. L. (2018). Quantitative epigenetics and evolution. Heredity, 121(3), 210–224. doi:10.1038/s41437-018-0114-x

Bates, D., Mächler, M., Bolker, B., & Walker, S. (2015). Fitting linear mixed-effects models using *lme4*. Journal of Statistical Software, 67(1). doi:10.18637/jss.v067.i01

Benjamini, Y., & Hochberg, Y. (1995). Controlling the false discovery rate: a practical and powerful approach to multiple testing. Journal of the Royal Statistical Society: series B (Methodological), 57(1), 289–300. doi:10.1111/j.2517-6161.1995.tb02031.x

Bewick, A. J., & Schmitz, R. J. (2017). Gene body DNA methylation in plants. Current Opinion in Plant Biology, 36, 103–110. doi:10.1016/j.pbi.2016.12.007

Blanca, G., Cabezudo, B., Cueto, M., Fernández López, C., & Morales Torres, C. (2009). Flora Vascular de Andalucía Oriental. Volumen 3: Rosaceae–Lentibulariaceae: Consejería de Medio Ambiente, Junta de Andalucía, Sevilla.

Bolger, A. M., Lohse, M., & Usadel, B. (2014). Trimmomatic: a flexible trimmer for Illumina sequence data. Bioinformatics, 30(15), 2114–2120. doi:10.1093/bioinformatics/btu170

Boquete, M. T., Muyle, A., & Alonso, C. (2021). Plant epigenetics: phenotypic and functional diversity beyond the DNA sequence. American Journal of Botany, 108(4), 553–558. doi:10.1002/ajb2.1645

Bossdorf, O., Richards, C. L., & Pigliucci, M. (2008). Epigenetics for ecologists. Ecology Letters, 11(2), 106–115. doi:10.1111/j.1461-0248.2007.01130.x

Bouyer, D., Kramdi, A., Kassam, M., Heese, M., Schnittger, A., Roudier, F., & Colot, V. (2017). DNA methylation dynamics during early plant life. Genome Biology, 18(1), 179. doi:10.1186/s13059-017-1313-0

Chen, X., Ge, X., Wang, J., Tan, C., King, G. J., & Liu, K. (2015). Genome-wide DNA methylation profiling by modified reduced representation bisulfite sequencing in *Brassica rapa* suggests that epigenetic modifications play a key role in polyploid genome evolution. Frontiers in Plant Science, 6, 836. doi:10.3389/fpls.2015.00836

Cokus, S. J., Feng, S., Zhang, X., Chen, Z., Merriman, B., Haudenschild, C. D., Pradhan, S., Nelson, S. F., & Pellegrini, M., & Jacobsen, S. E. (2008). Shotgun bisulphite sequencing of the *Arabidopsis* genome reveals DNA methylation patterning. Nature, 452(7184), 215–219. doi:10.1038/nature06745

Conesa, A., Madrigal, P., Tarazona, S., Gomez-Cabrero, D., Cervera, A., McPherson, A., Szcześniak, M. W., Gaffney, D. J., Elo, L. L., Zhang, X., & Mortazavi, A. (2016). A survey of best practices for RNA-seq data analysis. Genome Biology, 17, 13. doi:10.1186/s13059-016-0881-8

De Kort, H., Panis, B., Deforce, D., Van Nieuwerburgh, F., & Honnay, O. (2020). Ecological divergence of wild strawberry DNA methylation patterns at distinct spatial scales. Molecular Ecology, 29(24), 4871–4881. doi:10.1111/mec.15689

De Kort, H., Toivainen, T., Van Nieuwerburgh, F., Andres, J., Hytonen, T. P., & Honnay, O. (2022). Signatures of polygenic adaptation align with genome-wide methylation patterns in wild strawberry plants. New Phytologist, 235(4), 1501–1514. doi:10.1111/nph.18225

Denkena, J., Johannes, F., & Colome-Tatche, M. (2021). Region-level epimutation rates in *Arabidopsis thaliana*. Heredity, 127(2), 190–202. doi:10.1038/s41437-021-00441-w

Du, J., Zhong, X., Bernatavichute, Y. V., Stroud, H., Feng, S., Caro, E., Vashisht, A. A., Terragni, J., Chin, H. G., Tu, A., Hetzel, J., Wohlschlegel, J. A., Pradhan, S., Patel, D.J., & Jacobsen, S. E. (2012). Dual binding of chromomethylase domains to H3K9me2-containing nucleosomes directs DNA methylation in plants. Cell, 151(1), 167–180. doi:10.1016/j.cell.2012.07.034

Dubin, M. J., Zhang, P., Meng, D., Remigereau, M. S., Osborne, E. J., Paolo Casale, F., Drewe, P., Kahles, A., Jean, G., Vilhjálmsson, B., Jagoda, J., Irez, S., Voronin, V., Song, Q., Long, Q., Rätsch, G., Stegle, O., Clark, R. M., & Nordborg, M. (2015). DNA methylation in *Arabidopsis* has a genetic basis and shows evidence of local adaptation. eLife, 4, e05255. doi:10.7554/eLife.05255

Feng, H., Conneely, K. N., & Wu, H. (2014). A Bayesian hierarchical model to detect differentially methylated loci from single nucleotide resolution sequencing data. Nucleic Acids Research, 42(8), e69. doi:10.1093/nar/gku154

Fiz-Palacios, O., Vargas, P., Vila, R., Papadopulos, A. S., & Aldasoro, J. J. (2010). The uneven phylogeny and biogeography of *Erodium* (Geraniaceae): radiations in the Mediterranean and recent recurrent intercontinental colonization. Annals of Botany 106: 871–884. doi:10.1093/aob/mcq184

Galanti, D., Ramos-Cruz, D., Nunn, A., Rodríguez-Arevalo, I., Scheepens, J. F., Becker, C., & Bossdorf, O. (2022). Genetic and environmental drivers of large-scale epigenetic variation in *Thlaspi arvense*. PLoS Genetics, 18(10), e1010452. doi:10.1371/journal.pgen.1010452

Gawehns, F., Postuma, M., van Antro, M., Nunn, A., Sepers, B., Fatma, S., van Gurp, T. P., Wagemaker, N. C. A. M., Mateman, A. C., Milanovic-Ivanovic, S., Groβe, I., van Oers, K., Vergeer, P., & Verhoeven, K. J. F. (2022). epiGBS2: Improvements and evaluation of highly multiplexed, epiGBS-based reduced representation bisulfite sequencing. Molecular Ecology Resources, 22(5), 2087–2104. doi:10.1111/1755-0998.13597

Hawes, N. A., Fidler, A. E., Tremblay, L. A., Pochon, X., Dunphy, B. J., & Smith, K. F. (2018). Understanding the role of DNA methylation in successful biological invasions: a review. Biological Invasions, 20(9), 2285–2300. doi:10.1007/s10530-018-1703-6

Herrera, C. M., Alonso, C., Medrano, M., Perez, R., & Bazaga, P. (2018). Transgenerational epigenetics: Inheritance of global cytosine methylation and methylation-related epigenetic markers in the shrub *Lavandula latifolia*. American Journal of Botany, 105(4), 741–748. doi:10.1002/ajb2.1074

Herrera, C. M., Medrano, M., Bazaga, P., & Alonso, C. (2022). Ecological significance of intraplant variation: Epigenetic mosaicism in *Lavandula latifolia* plants predicts extant and transgenerational variability of fecundity-related traits. Journal of Ecology, 110(11), 2555–2567. doi:10.1111/1365-2745.13964

Hsu, F. M., Yen, M. R., Wang, C. T., Lin, C. Y., Wang, C. R., & Chen, P. Y. (2017). Optimized reduced representation bisulfite sequencing reveals tissue-specific mCHH islands in maize. Epigenetics Chromatin, 10(1), 42. doi:10.1186/s13072-017-0148-y

Ito, H., Gaubert, H., Bucher, E., Mirouze, M., Vaillant, I., & Paszkowski, J. (2011). An siRNA pathway prevents transgenerational retrotransposition in plants subjected to stress. Nature, 472(7341), 115–119. doi:10.1038/nature09861

Ito, H., Kim, J. M., Matsunaga, W., Saze, H., Matsui, A., Endo, T. A., Harukawa Y., Takagi H., Yaegashi H., Masuta Y., Masuda S., Ishida J., Tanaka M., Takahashi S., Morosawa T., Toyoda T., Kakutani T., Kato A., & Seki, M. (2016). A stress-activated transposon in *Arabidopsis* induces transgenerational abscisic acid insensitivity. Scientific Reports, 6, 23181. doi:10.1038/srep23181

Kidwell, M. G. (2002). Transposable elements and the evolution of genome size in eukaryotes. Genetica, 115, 49–63. doi:10.1023/a:1016072014259.

Klein, S. P., & Anderson, S. N. (2022). The evolution and function of transposons in epigenetic regulation in response to the environment. Current Opinion in Plant Biology, 69, 102277. doi:10.1016/j.pbi.2022.102277

Kolde, R. (2019). pheatmap: pretty heatmaps. R package version 1.0.12. Retrieved from https://CRAN.R-project.org/package=pheatmap

Korneliussen, T. S., Albrechtsen, A., & Nielsen, R. (2014). ANGSD: Analysis of Next Generation Sequencing Data. BMC Bioinformatics, 15(356). doi:10.1186/s12859-014-0356-4

Kreutz, C., Can, N. S., Bruening, R. S., Meyberg, R., Merai, Z., Fernandez-Pozo, N., & Rensing, S. A. (2020). A blind and independent benchmark study for detecting differeally methylated regions in plants. Bioinformatics, 36(11), 3314–3321. doi:10.1093/bioinformatics/btaa191

Krueger, F. (2013). Reduced Representation Bisulfite-Seq – A Brief Guide to RRBS. Retrieved from https://www.bioinformatics.babraham.ac.uk/projects/bismark/RRBS_Guide.pdf

Krueger, F., & Andrews, S. R. (2011). Bismark: a flexible aligner and methylation caller for Bisulfite-Seq applications. Bioinformatics, 27(11), 1571–1572. doi:10.1093/bioinformatics/btr167

Lamka, G. F., Harder, A. M., Sundaram, M., Schwartz, T. S., Christie, M. R., DeWoody, J. A., & Willoughby, J. R. (2022). Epigenetics in ecology, evolution, and conservation. Frontiers in Ecology and Evolution, 10, 871791. doi:10.3389/fevo.2022.871791

Langmead, B., & Salzberg, S. L. (2012). Fast gapped-read alignment with Bowtie 2. Nature Methods, 9(4), 357–359. doi:10.1038/nmeth.1923

Law, J. A., & Jacobsen, S. E. (2010). Establishing, maintaining and modifying DNA methylation patterns in plants and animals. Nature Reviews Genetics, 11(3), 204–220. doi:10.1038/nrg2719

Lea, A. J., Vilgalys, T. P., Durst, P. A. P., & Tung, J. (2017). Maximizing ecological and evolutionary insight in bisulfite sequencing data sets. Nature Ecology & Evolution, 1(8), 1074–1083. doi:10.1038/s41559-017-0229-0

Lenth, R. (2023). emmeans: Estimated marginal means, aka least-squares means. Retrieved from https://CRAN.R-project.org/package=emmeans

Matzke, M. A., & Mosher, R. A. (2014). RNA-directed DNA methylation: an epigenetic pathway of increasing complexity. Nature Reviews Genetics, 15(6), 394–408. doi:10.1038/nrg3683

Medrano, M., Alonso, C., Bazaga, P., López, E., & Herrera, C. M. (2020). Comparative genetic and epigenetic diversity in pairs of sympatric, closely related plants with contrasting distribution ranges in south-eastern Iberian mountains. AoB Plants, 12(3), plaa013. doi:10.1093/aobpla/plaa013

Moreno, J. C. (2008). Lista Roja 2008 de la Flora Vascular Española: Ministerio de Medio Ambiente y Medio Rural y Marino.

Muyle, A. M., Seymour, D. K., Lv, Y., Huettel, B., & Gaut, B. S. (2022). Gene body methylation in plants: Mechanisms, functions, and important implications for understanding evolutionary processes. Genome Biology and Evolution, 14(4). doi:10.1093/gbe/evac038

Nazareno, A. G., Bemmels, J. B., Dick, C. W., & Lohmann, L. G. (2017). Minimum sample sizes for population genomics: an empirical study from an Amazonian plant species. Molecular Ecology Resources, 17(6), 1136–1147. doi:10.1111/1755-0998.12654

Noshay, J. M., & Springer, N. M. (2021). Stories that can’t be told by SNPs; DNA methylation variation in plant populations. Current Opinion in Plant Biology, 61, 101989. doi:10.1016/j.pbi.2020.101989

Nunn, A., Can, S. N., Otto, C., Fasold, M., Díez Rodríguez, B., Fernández-Pozo, N., Rensing, S. A., Stadler, P. F., & Langenberger, D. (2021). EpiDiverse Toolkit: a pipeline suite for the analysis of bisulfite sequencing data in ecological plant epigenetics. NAR Genomomics Bioinformatics, 3(4), lqab106. doi:10.1093/nargab/lqab106

Oksanen, J., Simpson, G. L., Blanchet, F. G., Kindt, R., Legendre, P., Minchin, P. R., O’Hara, R. B., Simpson, G. L., Solymos, P., Stevens, M., Szoecs, E., Wagner, H., Barbour, M., Bedward, M., Bolker, B., Borcard, D., Carvalho, G., Chirico, M., De Caceres, M., Durand, S., & Weedon, J. (2022). Vegan: Community ecology package. R package version 2.6-2. Retrieved from https://CRAN.R-project.org/package=vegan

Park, Y., & Wu, H. (2016). Differential methylation analysis for BS-seq data under general experimental design. Bioinformatics, 32(10), 1446–1453. doi:10.1093/bioinformatics/btw026

Paun, O., Verhoeven, K. J. F., & Richards, C. L. (2019). Opportunities and limitations of reduced representation bisulfite sequencing in plant ecological epigenomics. New Phytologist, 221(2), 738–742. doi:10.1111/nph.15388

Purcell, S., Neale, B., Todd-Brown, K., Thomas, L., Ferreira, M. A., Bender, D., Maller, J., Sklar, P., De Bakker, P. I., Daly, M. J., & Sham, P. C. (2007). PLINK: a tool set for whole-genome association and population-based linkage analyses. American Journal of Human Genetics, 81(3), 559–575. doi:10.1086/519795

Pustahija, F., Brown, S. C., Bogunić, F., Bašić, N., Muratović, E., Ollier, S., Hidalgo, O., Bourge, M., Stevanović, V., & Siljak-Yakovlev, S. (2013). Small genomes dominate in plants growing on serpentine soils in West Balkans, an exhaustive study of 8 habitats covering 308 taxa. Plant and Soil, 373(1-2), 427–453. doi:10.1007/s11104-013-1794-x

R Core Team (2023). R: A language and environment for statistical computing. R Foundation for Statistical Computing.

Renaud, G., Stenzel, U., Maricic, T., Wiebe, V., & Kelso, J. (2015). deML: robust demultiplexing of Illumina sequences using a likelihood-based approach. Bioinformatics, 31(5), 770–772. doi:10.1093/bioinformatics/btu719

Richards, C. L., Alonso, C., Becker, C., Bossdorf, O., Bucher, E., Colomé-Tatché, M., Durka, W., Engelhardt, J., Gaspar, B., Gogol-Döring, A., Grosse, I., van Gurp, T. P., Heer, K., Kronholm, I., Lampei, C., Latzel, V., Mirouze, M., Opgenoorth, L., Paun, O., & Verhoeven, K. J. F. (2017). Ecological plant epigenetics: Evidence from model and non-model species, and the way forward. Ecology Letters, 20(12), 1576–1590. doi:10.1111/ele.12858

Richards, C. L., Bossdorf, O., & Verhoeven, K. J. F. (2010). Understanding natural epigenetic variation. New Phytologist, 187(3), 562–564.

Richards, E. J. (2006). Inherited epigenetic variation —revisiting soft inheritance. Nature Reviews, 7, 395–401.

Rochette, N. C., & Catchen, J. M. (2017). Deriving genotypes from RAD-seq short-read data using Stacks. Nature Protocols, 12(12), 2640–2659. doi:10.1038/nprot.2017.123

Schmitz, R. J., Schultz, M. D., Lewsey, M. G., O’Malley, R. C., Urich, M. A., Libiger, O., Schork, N. J., & Ecker, J. R. (2011). Transgenerational epigenetic instability is a source of novel methylation variants. Science, 334, 369–373. doi: 10.1126/science.1212959

Skotte, L., Korneliussen, T. S., & Albrechtsen, A. (2013). Estimating individual admixture proportions from next generation sequencing data. Genetics, 195(3), 693–702. doi:10.1534/genetics.113.154138

Stroud, H., Do, T., Du, J., Zhong, X., Feng, S., Johnson, L., Patel, D. J., & Jacobsen, S. E. (2014). Non-CG methylation patterns shape the epigenetic landscape in *Arabidopsis*. Nature Structural and Molecular Biology, 21(1), 64–72. doi:10.1038/nsmb.2735

Szukala, A., Bertel, C., Frajman, B., Schonswetter, P., & Paun, O. (2023). Parallel adaptation to lower altitudes is associated with enhanced plasticity in *Heliosperma pusillum* (Caryophyllaceae). The Plant Journal, 115(6), 1619–1632. doi:10.1111/tpj.16342

Tan, G., Polychronopoulos, D., & Lenhard, B. (2019). CNEr: a toolkit for exploring extreme noncoding conservation. PLoS Computational Biology, 15(8), e1006940.

Trucchi, E., Mazzarella, A. B., Gilfillan, G. D., Lorenzo, M. T., Schonswetter, P., & Paun, O. (2016). BsRADseq: screening DNA methylation in natural populations of non-model species. Molecular Ecology, 25(8), 1697–1713. doi:10.1111/mec.13550

Valverde, J., Medrano, M., Herrera, C. M., & Alonso, C. (2024). Comparative epigenetic and genetic spatial structure in Mediterranean mountain plants: a multispecies study. Heredity, 132(2), 106–116. doi:10.1038/s41437-024-00668-3

van der Graaf, A., Wardenaar, R., Neumann, D. A., Taudt, A., Shaw, R. G., Jansen, R. C., Schmitz, R. J., Colomé-Tatché, M., & Johannes, F. (2015). Rate, spectrum, and evolutionary dynamics of spontaneous epimutations. Proceedings of the National Academy of Sciences of the United States of America, 112(21), 6676–6681. doi:10.1073/pnas.1424254112

Varotto, S., Krugman, T., Aiese Cigliano, R., Kashkush, K., Kondic-Spika, A., Aravanopoulos, F. A., Pradillo, M., Consiglio, F., Aversano, R., Pecinka, A., & Miladinovic, D. (2022). Exploitation of epigenetic variation of crop wild relatives for crop improvement and agrobiodiversity preservation. Theoretical and Applied Genetics, 135(11), 3987–4003. doi:10.1007/s00122-022-04122-y

Vidalis, A., Zivkovic, D., Wardenaar, R., Roquis, D., Tellier, A., & Johannes, F. (2016). Methylome evolution in plants. Genome Biology, 17(1), 264. doi:10.1186/s13059-016-1127-5

Weng, M. L., Becker, C., Hildebrandt, J., Neumann, M., Rutter, M. T., Shaw, R. G., Weigel, D., & Fenster, C. B. (2019). Fine-grained analysis of spontaneous mutation spectrum and frequency in *Arabidopsis thaliana*. Genetics, 211(2), 703–714. doi:10.1534/genetics.118.301721

Williams, B. R., Miller, A. J., & Edwards, C. E. (2023). How do threatened plant species with low genetic diversity respond to environmental stress? Insights from comparative conservation epigenomics and phenotypic plasticity. Molecular Ecology Resources. doi:10.1111/1755-0998.13897

Záveská, E., Maylandt, C., Paun, O., Bertel, C., Frajman, B., The STEPPE Consortium, & Schönswetter, P. (2019). Multiple auto- and allopolyploidisations marked the Pleistocene history of the widespread Eurasian steppe plant *Astragalus onobrychis* (Fabaceae). Molecular Phylogenetics and Evolution, 139. doi:10.1016/j.ympev.2019.106572

Zeng, W., & Mortazavi, A. (2012). Technical considerations for functional sequencing assays. Nature Immunology, 13(9), 802–807. doi:10.1038/ni.2407

Zhao, L., Xie, L., Zhang, Q., Ouyang, W., Deng, L., Guan, P., Ma, M., Li, Y., Zhang, Y., Xiao, Q., Zhang, J., Li, H., Wang, S., Man, J., Cao, Z., Zhang, Q., Zhang, Q., Li, G., & Li, X. (2020). Integrative analysis of reference epigenomes in 20 rice varieties. Nature Communications, 11(1), 2658. doi:10.1038/s41467-020-16457-5

Zonneveld, B. J. M. (2019). The DNA weights per nucleus (genome size) of more than 2350 species of the Flora of The Netherlands, of which 1370 are new to science, including the pattern of their DNA peaks. Forum geobotanicum, 8, 24–78. doi:10.3264/FG.2019.1022

