## Supplementary Information for "Decoupled genetic and epigenetic variation in the montane endemic *Erodium cazorlanum* (Geraniaceae) and a widespread congener"

#### Table of Contents:


| | |
|---|---|
| **RADseq library preparation** | Page 2 |
| **Genome assembly** | Page 3 |
| **Alternative population genomics analysis** | Page 6 |
| **Supplementary Figures** | Page 9 |
| **References** | Page 39 |

#### RADseq library preparation

Leaf DNA was digested with *Pst*I-HF restriction enzyme (New England Biolabs, R3140S) at 37 $^{o}$C for 1 h, followed by inhibition of the enzyme at 80 $^{o}$C for 20 min. Next, eight different P1 adapters were ligated to the digested DNA fragments using T4 ligase (New England Biolabs, M0202M) in a thermal cycler (incubation 16 $^{o}$C overnight, deactivation 65 $^{o}$C for 10 min). DNA samples were pooled into five P1 multiplexed mixes (containing DNA tagged with different P1 adapters), the pooled samples were fragmented with a Covaris with parameters *peak incident:175 duty factor: 10% cycles: 200 seconds: 8*, then cleaned with the MinElute Reaction Cleanup Kit (Qiagen 28204). DNA fragments were selected by size with SPRIselect magnetic beads (Beckman Coulter Life Sciences, B23318). DNA ends were phosphorylated for 30 min at room temperature with the Quick Blunt Kit (New England Biolabs, E1201S), ddATP was added with a Klenow exo enzyme (New England Biolabs, M0212S) at 37 $^{o}$C for 30 min. After DNA quantification, adenylated DNA ends from the P1-multiplexed pooled samples were ligated to P2 adapters with a T4 ligase (New England Biolabs, M0202M). Adapter combinations were unique to each sample (adapters listed in **Supplementary Table S13**). After ligation, pooled DNA samples were separated in two aliquots, one for RADseq and another for bsRADseq. Aliquots for bsRADseq were denatured at 50 $^{o}$C for 10 min, then treated with bisulfite prepared conversion reagent solution from the MethylEdge kit (Promega, N1301). Afterwards, aliquots were incubated in a thermal cycler with two cycles at 65 $^{o}$C 30 min, 95 $^{o}$C 90 sec, plus a cycle at 65 $^{o}$C 30 min, 4 $^{o}$C 30 min. After desalting (and desulfonating when needed) the samples, polymerase chain reaction (PCR) was performed with the KAPA HiFi Hotstart Uracil+ kit (Kapa, KK280), following manufacturer indications.

### Genome assembly

*DNA Sequencing*

DNA was isolated from leaves of a single *E. cicutarium* individual reared after two events of autogamous selfing under greenhouse conditions, using a high-molecular weight DNA isolation protocol (Schalamun & Schwessinger, 2017). Illumina and PacBio libraries preparation and sequencing were performed in AllGenetics (www.allgenetics.eu). HiSeq X Reagent Kit v2.5 (Illumina) was used to prepare paired end short reads library, and SMRTbell Express Template Prep Kit 2.0 (PacBio) was used to prepare the long reads library, following the manufacturer's instructions. The libraries were sequenced in a HiSeq X PE150 (Illumina) with an Illumina patterned flow cell, and a Sequel II sequencer (PacBio), with an SMRT Cell 8M under the Long-reads mode, respectively. Reads were quality-checked using the software FastQC v0.11.5 (Andrews, 2010) for Illumina reads and SequelTools (Hufnagel et al., 2020) for PacBio reads. Whole-genome sequencing in the Illumina HiSeq X PE150 platform yielded a total of 333,195,590 paired-end reads; and the PacBio Sequel II platform yielded a total of 6,757,035 subreads. **Supplementary Table S14** shows the sequencing statistics for both read sets.

*Genome de novo assembly*

Short and long reads we assembled *de novo* into "mega-reads" using the software MaSuRCA v3.4.2 (Zimin et al., 2017), with the short reads being error-corrected using QuorUM (Marcais et al., 2015). The assembler extends the short reads into "super-reads" (Zimin et al., 2013), which are then aligned to the long reads. Consistent alignments of "super-reads" are merged into "mega-reads". Finally, the "mega-reads" are assembled using the algorithm Flye v2.5 (Kolmogorov et al., 2019) implemented in MaSuRCA. A second *de novo* hybrid assembly was performed using the software HASLR (Haghshenas et al., 2020). The assemblies generated by MaSuRCA and HASLR were subsequently polished using POLCA (Zimin & Salzberg, 2020). Quality check of these assemblies was performed with QUAST 5.0.2 (Gurevich et al., 2013). The program Jellyfish (Marcais & Kingsford, 2011) was used to count the occurrences of kmers in the corrected reads setting the kmer size parameter to k = 31. GenomeScope (Vurture et al., 2017) was used to plot the k-mer profile **Supplementary Figure S18**. The number of scaffolds and the total length (in base pairs, bp) of the genome assembly, are recorded in **Supplementary Table S15**.

*Quality check of hybrid assembly*

The short sequencing reads were mapped to the genome assembly generated by MaSuRCA using the software BWA v0.7.15 (Li, 2013), in order to determine the coverage of each genomic region. SAMtools 1.3.1 (Li et al., 2009) calculated that 99.72% of the reads were mapped back to the assembly. BlobTools v1.1.1 was used to detect potential contaminant sequences in the assembly **Supplementary Figure S19**. The quality and completeness of the genome assembly was evaluated using BUSCO V5.beta.1 (Kollmar, 2019), setting the "genome" mode. The *Metaeuk* pipeline (Fadiji & Babalola, 2020) implemented in BUSCO was used to predict eukaryotic genes with default parameters, which were run with the lineage-specific *embryophyta_odb10* (last updated 2020-09-10) **Supplementary Figure S20**.

*Repetitive elements*

Repetitive elements were identified *de novo* in the assembly using RepeatModeler v2.0.1 (Flynn et al., 2020). Then, RepeatMasker v4.1.2-p1 (Smit et al., 2021) and the library of known repeats Repbase-20170127, were used to mask repeated elements from the assembly. A total of 329,448,982 bps (41.97 % of the assembly) was masked as repeated elements **Supplementary Table S16**.

*Genome annotation*

Gene prediction of the resulting assembly was performed using AUGUSTUS v3.2.3 (Stanke et al., 2006) by setting 'species = *Arabidopsis thaliana*' and 'gff = on'. The resulting prediction was analyzed with gffread v0.12.6 (Pertea & Pertea, 2020), which extracts the sequence of all transfrags (defined as transcripts or transcript fragments that result from the assembly process) and generates a FASTA file with these sequences. We then used TransDecoder v5.5.0 to identify candidate coding regions within mRNA sequences.

The predicted protein-coding genes were subsequently functionally annotated using InterProScan v5.50-84.0 (Jones et al., 2014). The annotation was performed against several general-content databases and included: the conserved domain database (CDD) (Lu et al., 2020), the Coils database (Lupas et al., 1991), the Gene3D database (Lewis et al., 2018), the HAMAP database (Pedruzzi et al., 2015), the MobiDBLite database (Necci et al., 2017), the protein analysis through evolutionary relationships (PANTHER) classification system (Mi et al., 2013), the protein families database (Pfam) (Finn et al., 2014), the protein information resource and superfamily (PIRSF) classification system (Wu et al., 2004), the protein motif fingerprints (PRINTS) database (Attwood et al., 1994), the protein domains, families and functional sites (ProSitePatterns and ProSiteProfiles) databases (Sigrist et al., 2013), the structure function linkage (SFLD) database (Akiva et al., 2014), the simple modular architecture research tools (SMART) (Letunic & Bork, 2018), the SUPERFAMILY database (Gough et al., 2001), and the TIGRFAM database (Haft et al., 2013).

To gain more information about the biological function of these potentially genes, a second annotation was performed using the software Sma3s v2 (Muñoz-Mérida et al., 2014), and the complete manually annotated and reviewed Swiss-Prot database from

UniProtKB (Consortium, 2007). Sma3s reports a summary with different categories and the number of sequences belonging to each functional category. Sequence annotations also include both the most probable gene name and the most probable description (including putative EC enzyme codes).

PacBio long reads are deposited in the sequence read archive (SRA) of the NCBI, under BioProject ID PRJNA984161. The resulting draft genome assembly of *E. cicutarium* is available at NCBI Genome repository, under accession JBDFRG000000000.

### Alternative population genetics analysis

*SNP calling and visualization*

An alternative pipeline to *STACKs* was conducted to test the consistency of SNP detection and population structure analyses. *ANGSD* (Korneliussen et al., 2014) was run on the BAM files produced by *Bowtie2* (Langmead & Salzberg, 2012) (which aligned RADseq data against the *E. cicutarium* draft genome), to conduct SNP calling. We calculated genotype likelyhoods using the *SAMtools* model in *ANGSD*, filtering positions with MAF < 0.11 and with inconsistent calling in more than 50 % of the individuals. A genetic distance matrix was also generated using the *–doIBS* and *–makeMatrix* options. *ANGSD* reported 180,348 and 235,107 SNPs, which overlapped with 65 and 144 DMCs in *E. cazorlanum* and *E. cicutarium*, respectively.

PCA analysis was run on SNP data with the script "*pcangsd.py*" (Meisner & Albrechtsen, 2018) using the covariance matrix generated by *ANGSD*, and PCA biplots were generated with R v4.1.2 (R Core Team, 2023). PCA did not reveal population genetic structure in *E. cazorlanum*, but showed divergence of the PT population in *E. cicutarium* **Supplementary Figure S21**. A heatmap of the pairwise co-variance matrix also showed no structure for *E. cazorlanum*, and some limited structure in *E. cicutarium* **Supplementary Figure S21**.

Genetic and epigenetic distance matrices were compared through Mantel's test using *vegan* package in R (Oksanen et al., 2022). Mantel's test showed borderline statistical significance between genetic and epigenetic distance matrices in *E. cazorlanum* ($P = 0.045$), while no statistical significance was found for *E. cicutarium* ($P > 0.9$).

*Admixture analysis*

*NGSadmix* (Skotte et al., 2013) was then run iteratively to generate 10 best likelyhood estimates for K values from one to ten. After formatting the outputs in Excel, *pong* (Behr et al., 2016) was run to generate a bar plot with the K proportions assigned to each individual with the best (highest) likelihood. Evanno method (Evanno et al., 2005) was then used to estimate ΔK between different K values, adapting the ΔK function found in STRUCTURE (Pritchard et al., 2000) into the Excel formula:

"*=ABS(mean_GL_K[n-1] - 2*mean_GL_K[n] + mean_GL_K[n+1])/SD_GL_K[n]*"

where *mean_GL_K[n]* is the average genotype likelihood of the analyzed K, *mean_GL_K[n-1]* and *mean_GL_K[n+1]* are the average genotype likelihood of the K values preceding and following the K value analyzed, and *SD_GL_K[n]* is the standard deviation of the genotype likelihood of the analyzed K. Evanno's method shows maximum ΔK values for K = 2 for both species **Supplementary Figure S22**. However, while there were no local ΔK maximums in *E. cazorlanum*, a local ΔK maximum was found for K = 4 in *E. cicutarium*. The plots generated by *pong* **Supplementary Figure S22** show again no population-congruent clustering of the samples in *E. cazorlanum*, while separation of population PT is patent in *E. cicutarium*.

*Comparison with results of main genetic analysis*

The number of SNPs has increased compared to the former analysis, due to lowering the minimum number of individuals with a correctly reported position (from 80 % to 50 %) and probably due to in-build methodological differences between *STACKS* and *ANGSD*.

In addition, 7,805 SNPs from *E. cazorlanum* and 13,183 SNPs from *E. cicutarium* detected by *ANGSD* were also detected by *STACKs*, both values were greater than expected by chance (hypergeometric test, p-value < 0.001), indicating that both pipelines managed to recover a similar set of SNPs.

Genetic population structure trends remained similar to the former analysis: identical for *E. cicutarium*, but a lot more mixed in *E. cazorlanum*. This might be a consequence of a higher number of markers analyzed by *ANGSD*. Mantel's test still shows no significant correlation between epigenetic and genetic distances in both species (*E. cazorlanum*: Mantel's r = 0.107, P-value = 0.051; *E. cicutarium*: Mantel's r = -0.177, P-value = 0.934).

When Mantel's tests were run comparing the SNP genetic distance with the epigenetic distance calculated from DMCs from the different methylation contexts separately (CpG, CHG and CHH), all comparisons were non-significant (p-value > 0.05) with the exception of CHH in *E. cazorlanum* (Mantel's r = 0.121, p-value = 0.038).

Evanno's method proved that K = 2 was the most likely number of ancestral lineages for both species, but taking into account its limitations (i. e. inability to detect panmixia or cases where K is optimal at 1), and regarding PCA and the admixture bar plot, we could arguably interpret local maximum values of ΔK as the ones closer to being true (i. e., K = 4 for *E. cicutarium*). Following the trend, admixture analysis assigned the ancestral groups loosely among individuals in *E. cazorlanum*, and in a more population-driven fashion in *E. cicutarium*.

### Supplementary Figures

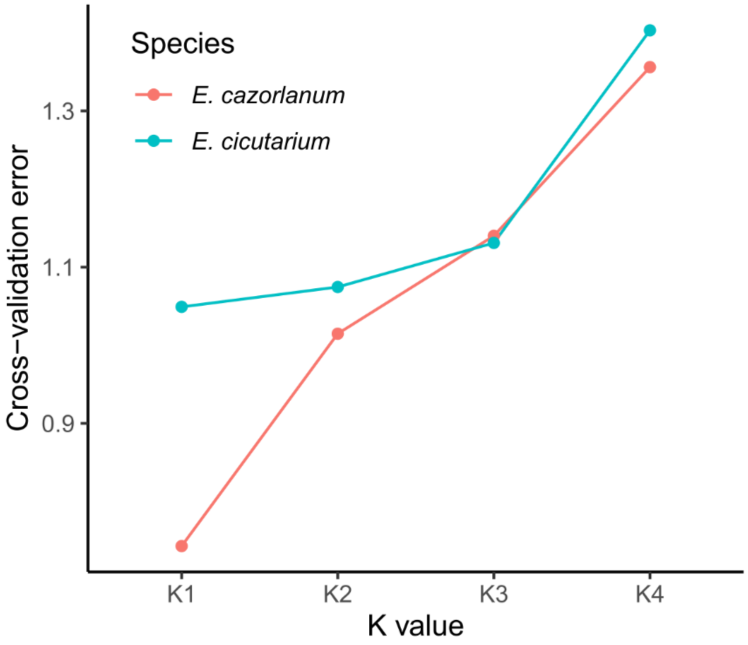


**Supplementary Figure S1.** Cross-validation error values from ADMIXTURE analysis. X axis shows the K values used for each tested model. Y axis shows the cross-validation error for each model. The red line shows *E. cazorlanum* values, and the blue line shows *E. cicutarium* values.

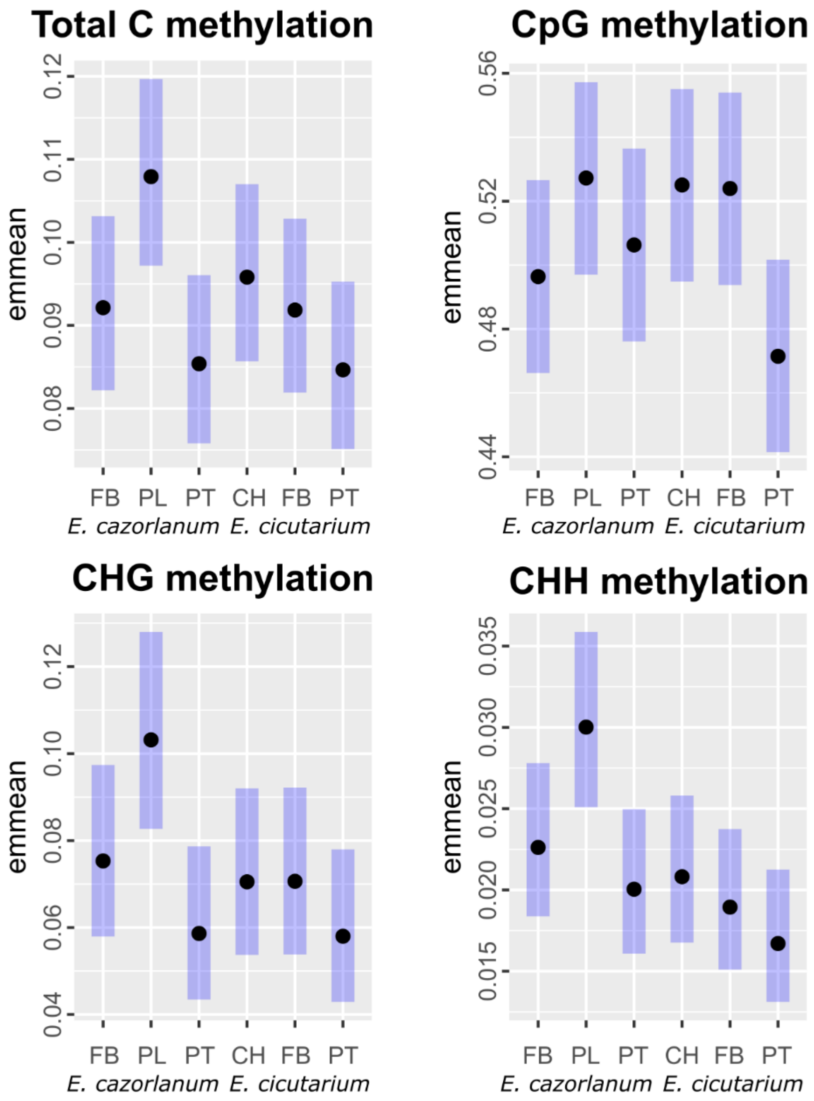


**Supplementary Figure S2.** Average relative proportions of methylated vs unmethylated cytosines in populations of the two *Erodium* species studied. Estimate marginal means (*emmeans*) for all the cytosines in the genome (top left) and for the methylation contexts CpG (top right), CHG (bottom left) and CHH (bottom right) are represented in each plot as black dots. X axis shows both *Erodium* species and populations, Y axis shows the estimated values for the proportion of methylated cytosines found by population obtained from quasibinomial GLM with species and populations included as fixed factors. Dispersion is shown as a shaded blue bar behind the dots for each population. All differences between species and populations were statistically not significant (P-value > 0.05).

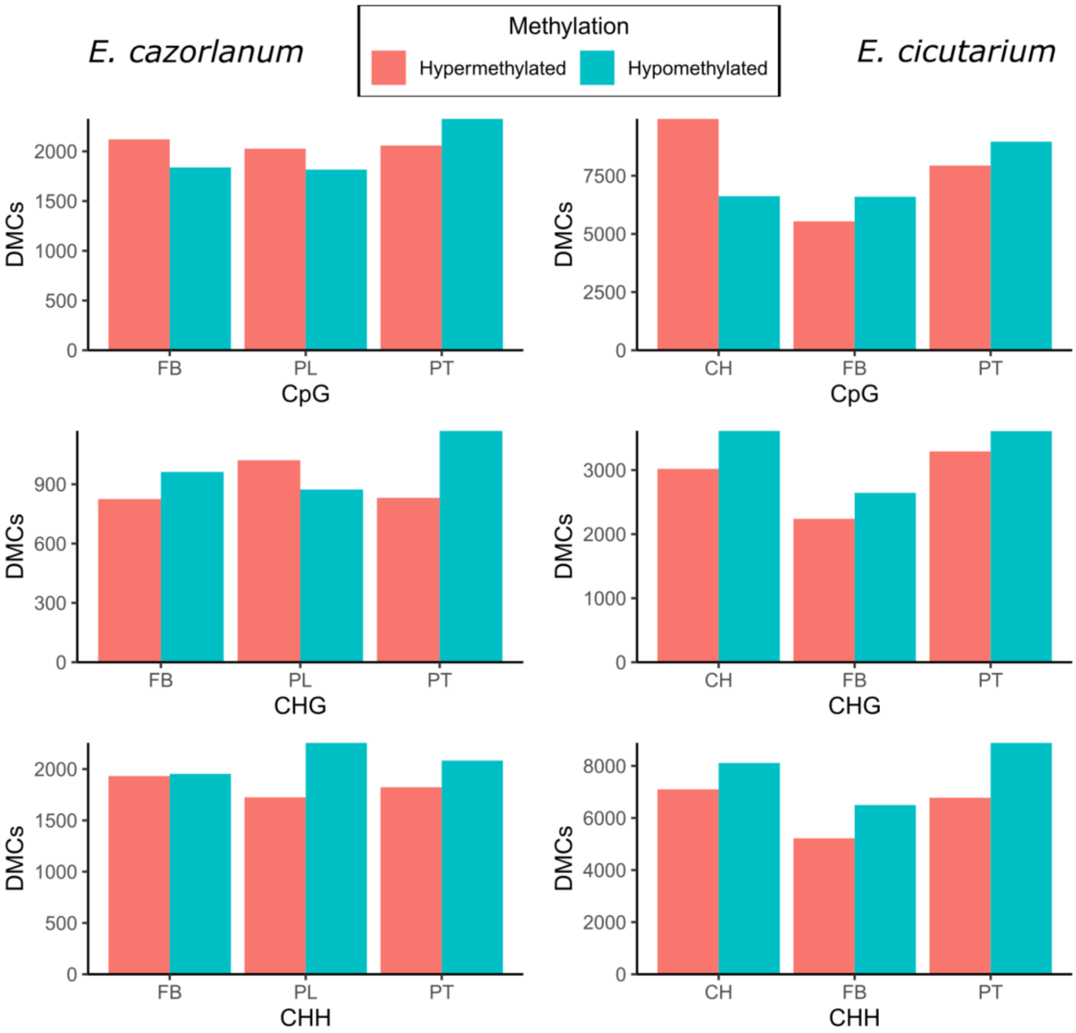


**Supplementary Figure S3.** Population specific differentially methylated cytosines (DMCs). Barplots show the number of hyper- (red) and hypo-methylated (blue) DMCs detected for each dataset. The left column shows *E. cazorlanum* DMCs, the right column shows *E. cicutarium* DMCs. X axis shows populations (FB = Fuente Bermejo, PL = Puerto Lézar, PT = Puerto del Tejo, CH = Nava Correhuelas), Y axis shows number of detected DMCs.

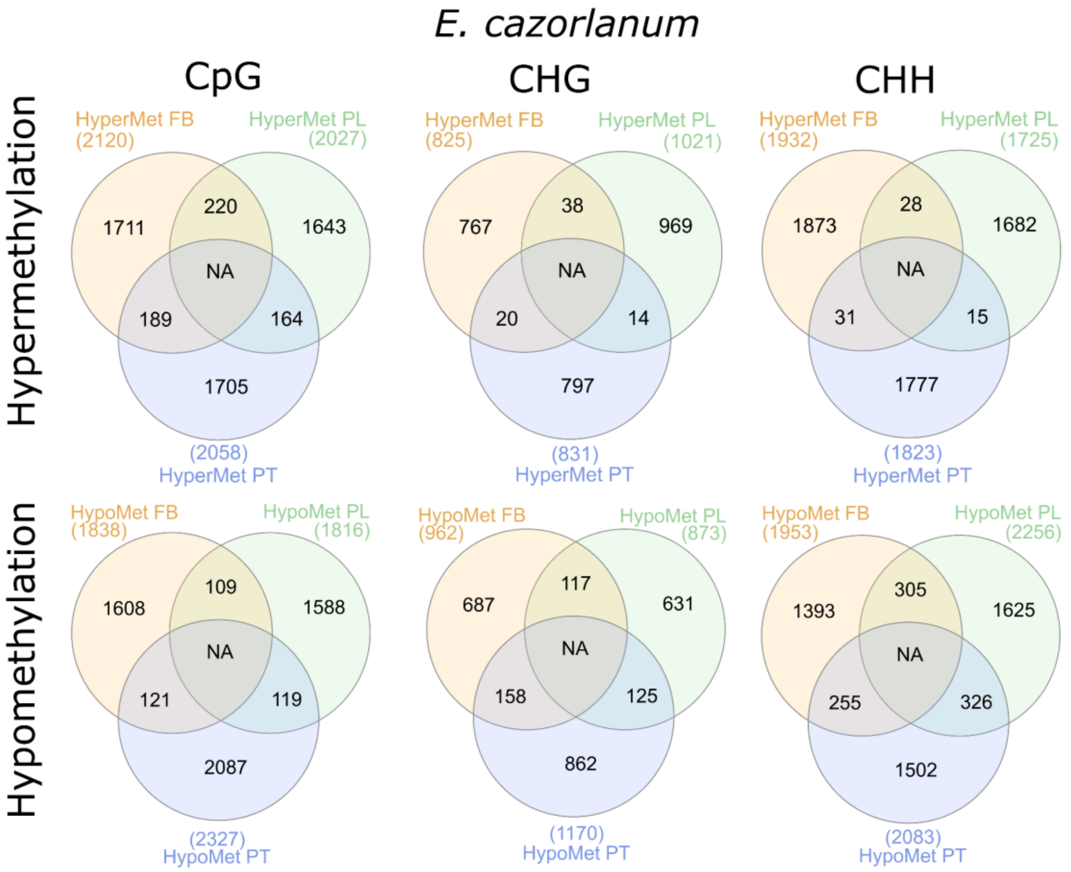


**Supplementary Figure S4.** Overlapping DMCs between *E. cazorlanum* populations. The top row shows values for hyper-methylated DMCs, bottom row shows the values for hypo-methylated DMCs. Each column shows a different methylation context. Venn diagrams compare the number of DMCs for the three different populations and their overlaps. Numbers in parentheses show the total number of DMCs per population.

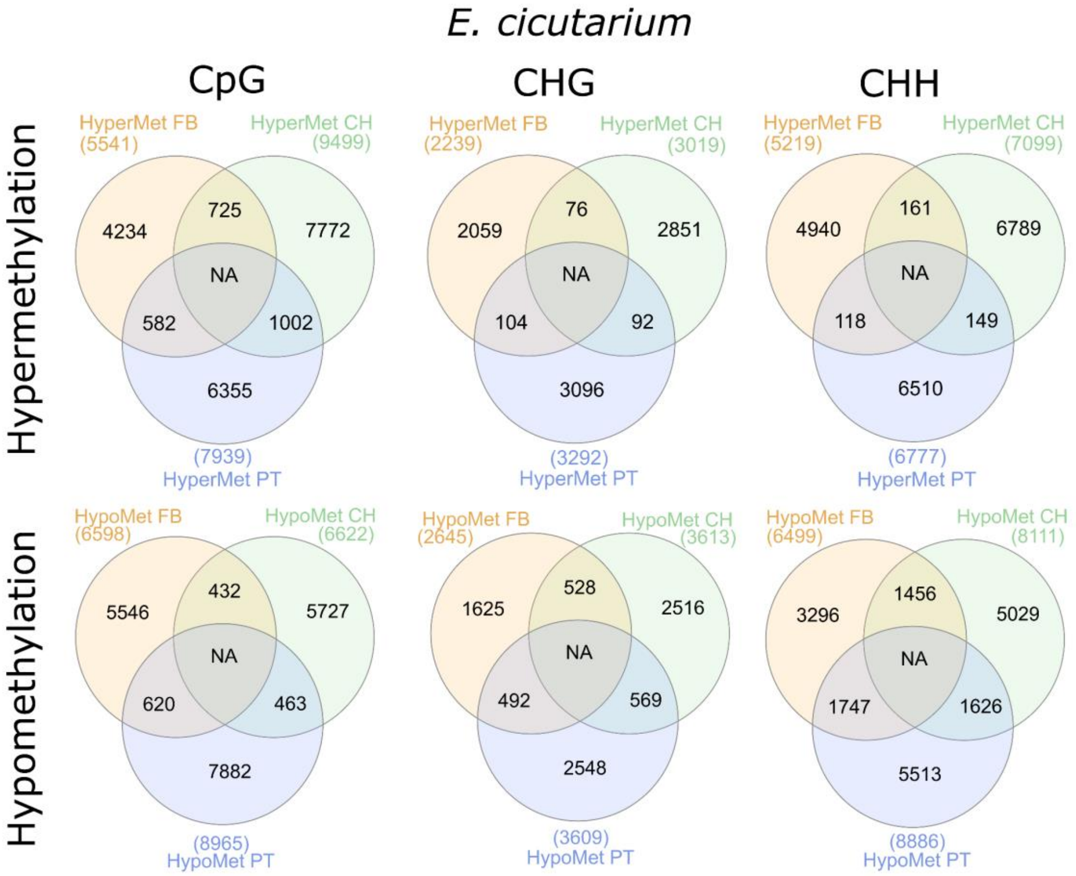


**Supplementary Figure S5.** Overlapping DMCs between *E. cicutarium* populations. The top row shows values for hyper-methylated DMCs, bottom row shows the values for hypo-methylated DMCs. Each column shows a different methylation context. Venn diagrams compare the number of DMCs for the three different populations and their overlaps. Numbers in parentheses show the total number of DMCs per population.

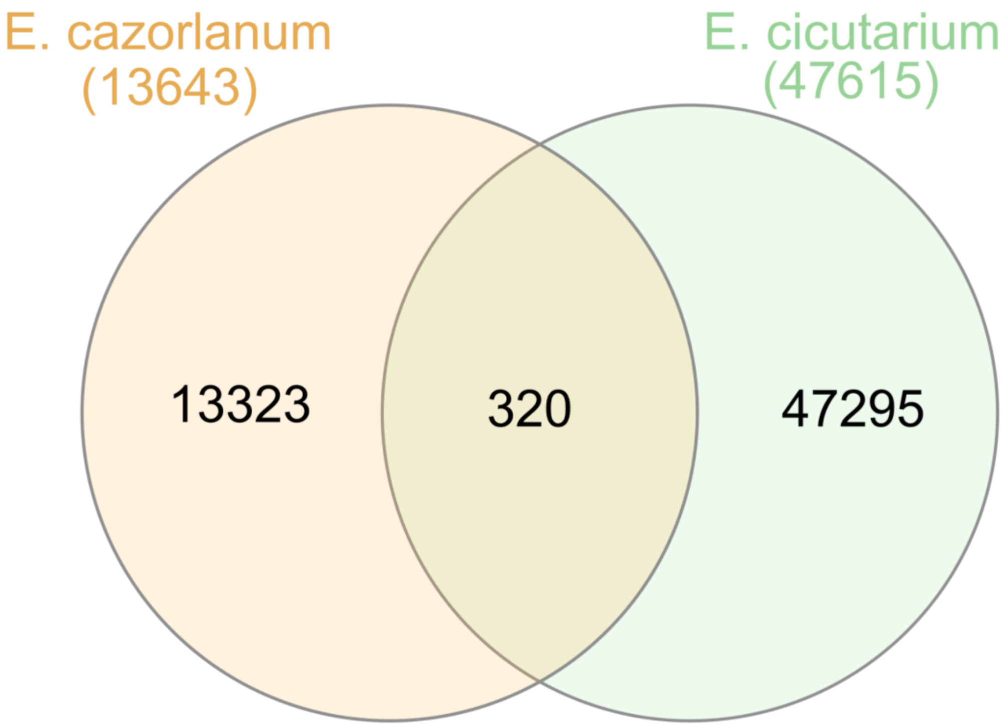


**Supplementary Figure S6.** Venn diagram with the number of DMCs detected between populations of each study species and those that were present in the two species. Numbers in parentheses show the total number of DMCs per species.

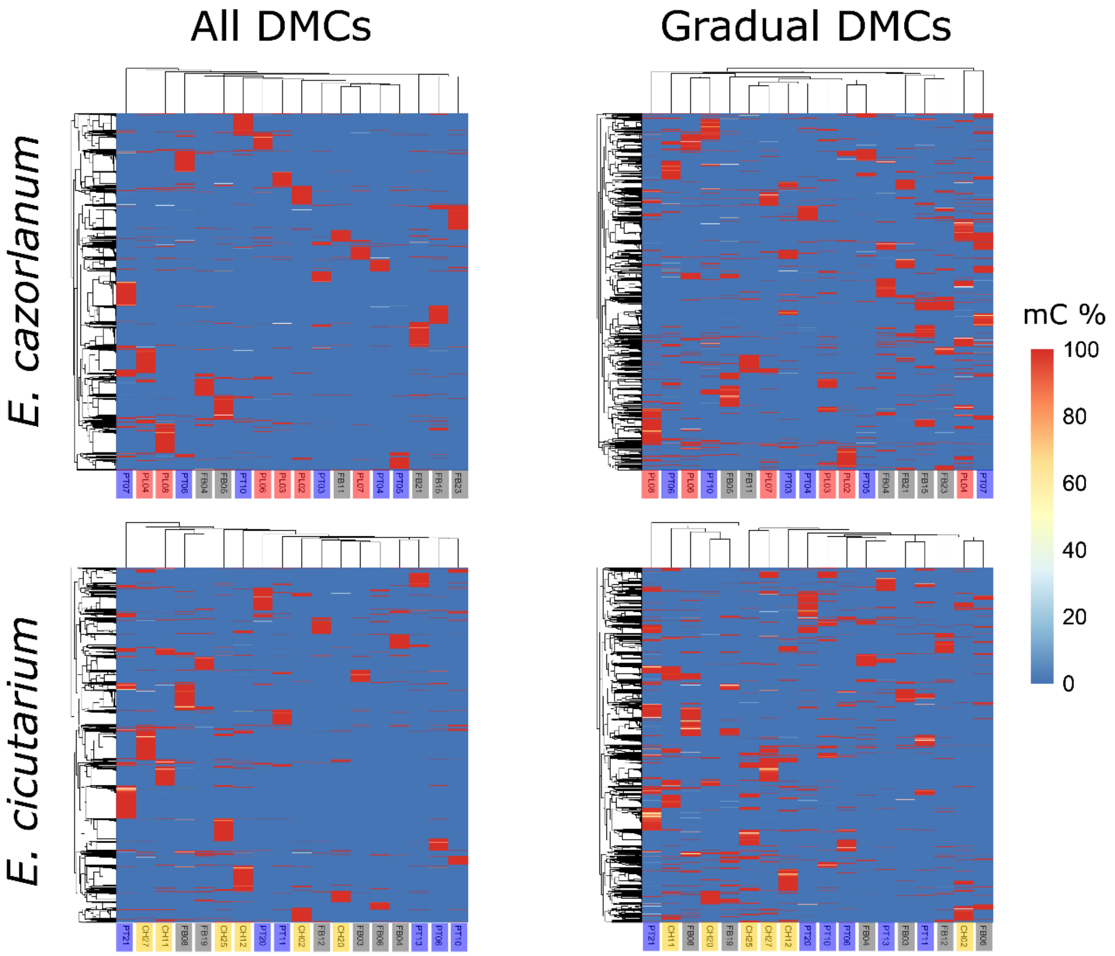


**Supplementary Figure S7.** Heatmap for DMC methylation percentage and hierarchical clustering of individuals based on percentage of methylation at detected DMCs. Top row shows data for *E. cazorlanum*, bottom row shows data for *E. cicutarium*. All DMCs and only DMCs with "gradual" methylation changes between samples (i.e., excluding DMC loci that were non-methylated in all except one individual which had ca. 100 % methylation) are shown in the left and the right column, respectively. X axis shows the identity of individual samples arranged by hierarchical clustering and colored according to population of origin. Y axis shows the different DMCs, arranged by hierarchical clustering. Colored tiles show the methylation percentage values per DMC and sample, ranging from 0 % (deep blue) to 100 % (red) methylation percentage.

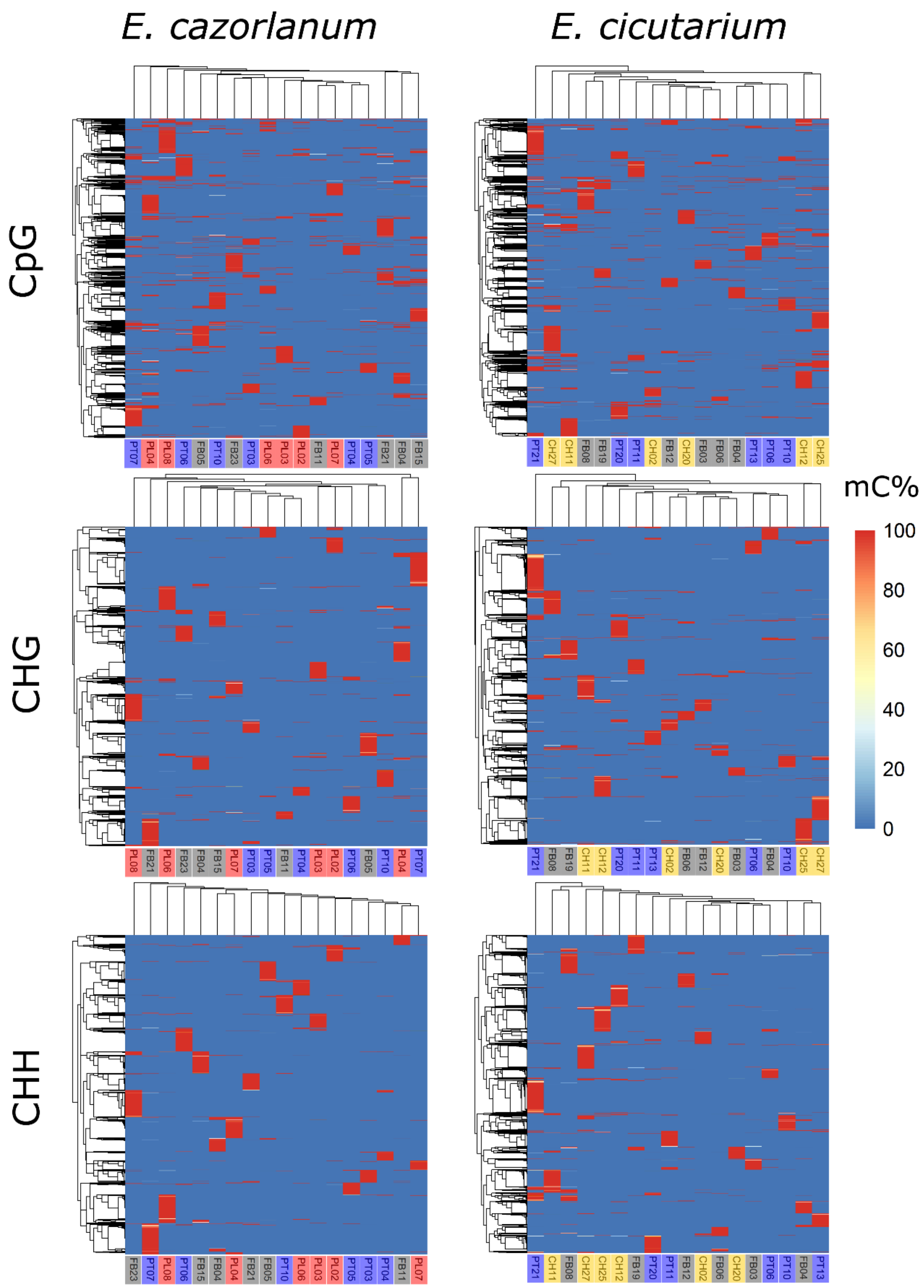


**Supplementary Figure S8.** Heatmap for DMC methylation percentage and hierarchical clustering of individuals based on percentage of methylation from all detected DMCs in each of the three different methylation contexts (CpG, CHG, and CHH). The left column shows the results for *E. cazorlanum* and the right column shows the results for *E. cicutarium*. Each row corresponds to CpG (top), CHG (middle), and CHH (bottom) methylation contexts. X axis shows the identity of different individual samples arranged by hierarchical clustering. Y axis shows the different DMCs, arranged by hierarchical clustering. Colored tiles show the methylation percentage values per DMC and sample, ranging from 0 % (deep blue) to 100 % (red) methylation percentage.

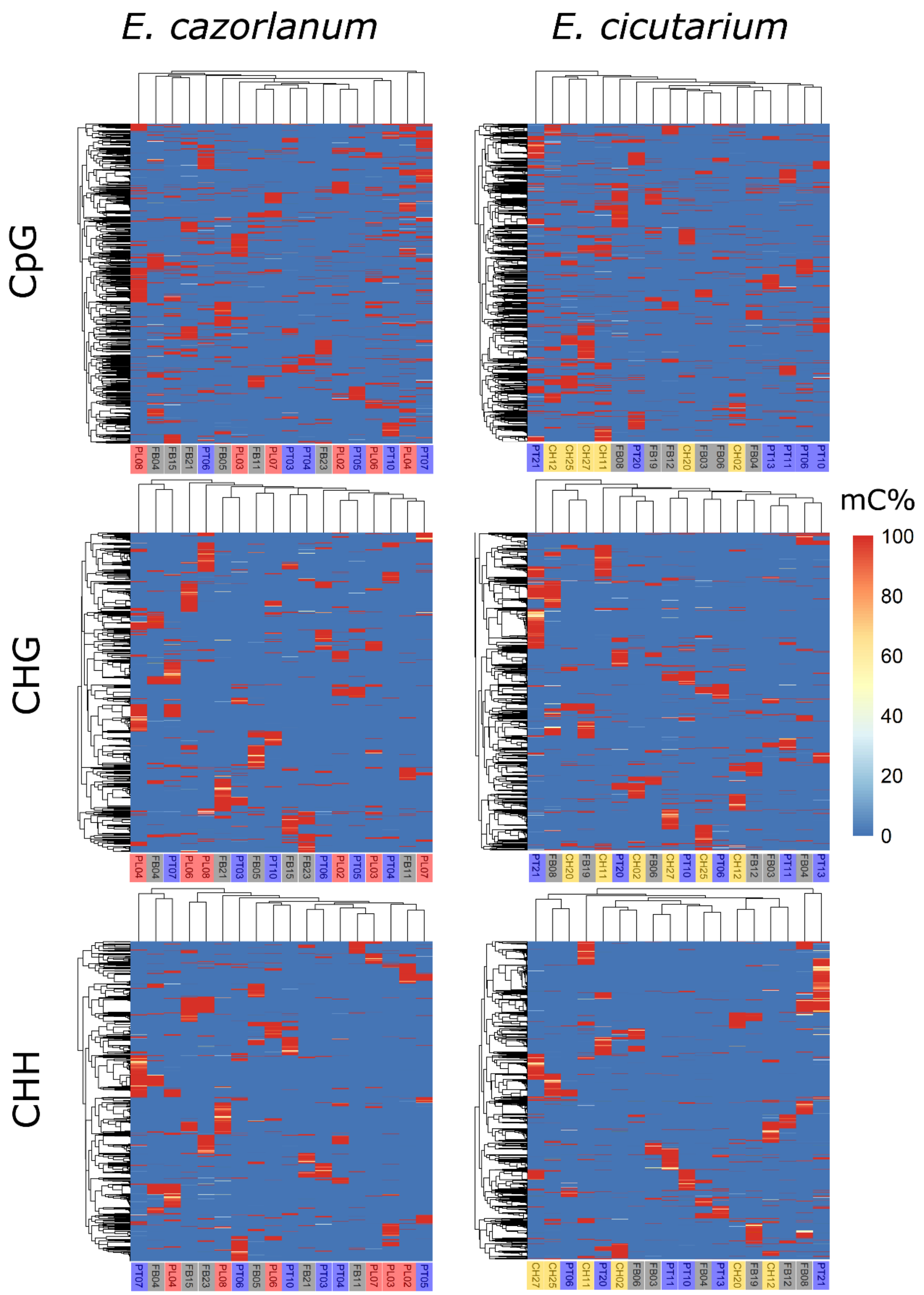


**Supplementary Figure S9.** Heatmap for DMC methylation percentage and hierarchical clustering of individuals based on percentage of methylation from gradual DMCs in each of the three different methylation contexts (CpG, CHG, and CHH). The left column shows the results for *E. cazorlanum* and the right column shows the results for *E. cicutarium*. Each row corresponds to CpG (top), CHG (middle), and CHH (bottom) methylation contexts. X axis shows the identity of different individual samples arranged by hierarchical clustering. Y axis shows the different DMCs, arranged by hierarchical clustering. Colored tiles show the methylation percentage values per DMC and sample, ranging from 0 % (deep blue) to 100 % (red) methylation percentage.

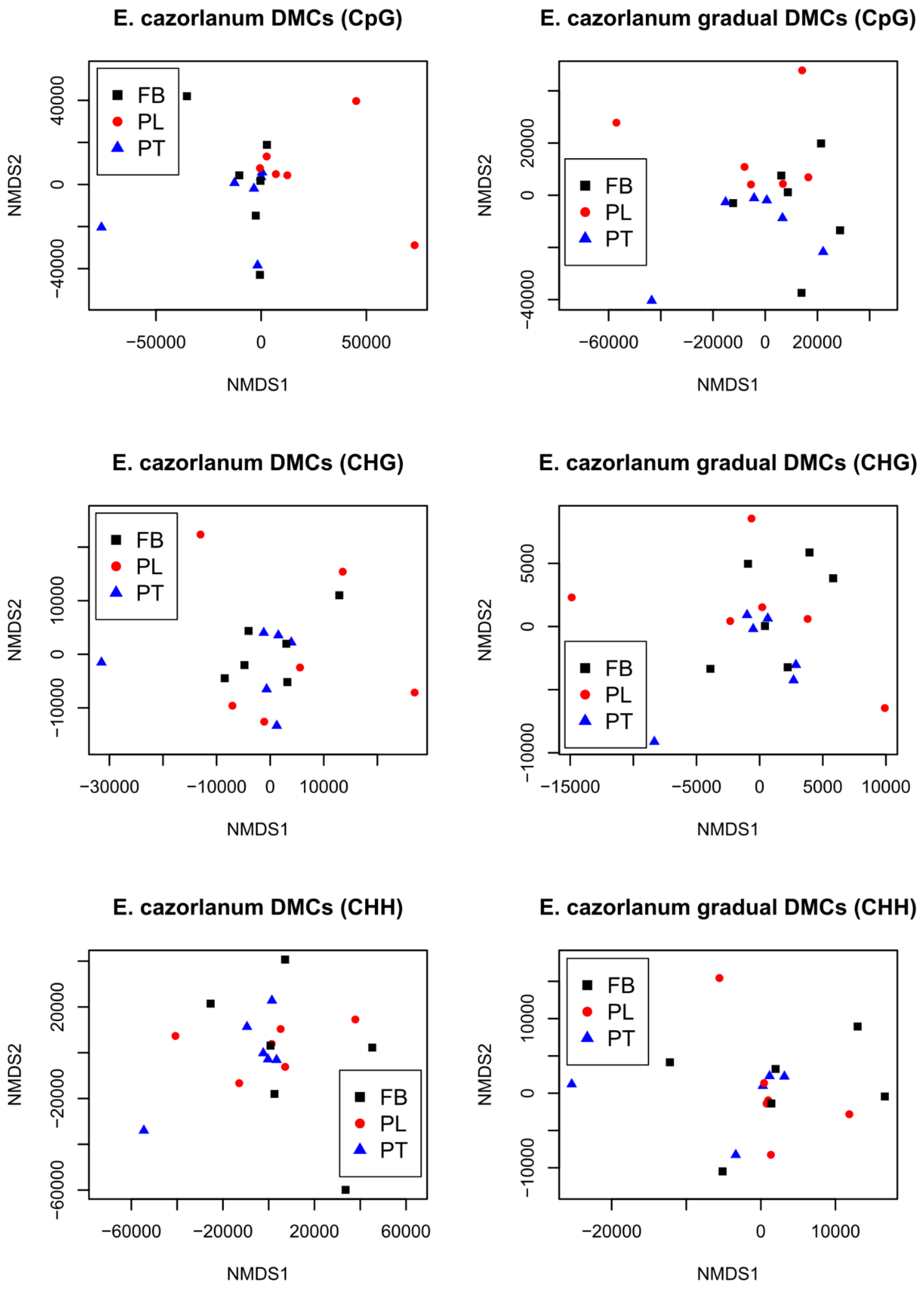


**Supplementary Figure S10.** Multilocus analysis of individual variation in methylation percentage of DMCs of *E. cazorlanum* populations separated by context. CpG, CHG and CHH are shown from top to bottom rows, respectively. Left column shows the output of non-metric multidimensional scaling (NMDS) for all the DMCs, while the right column shows the output for the gradual DMCs. X and Y axes show NMDS analysis dimensions 1 and 2, respectively. Colors of data points show the population of origin (black: FB, red: PL, blue: PT).

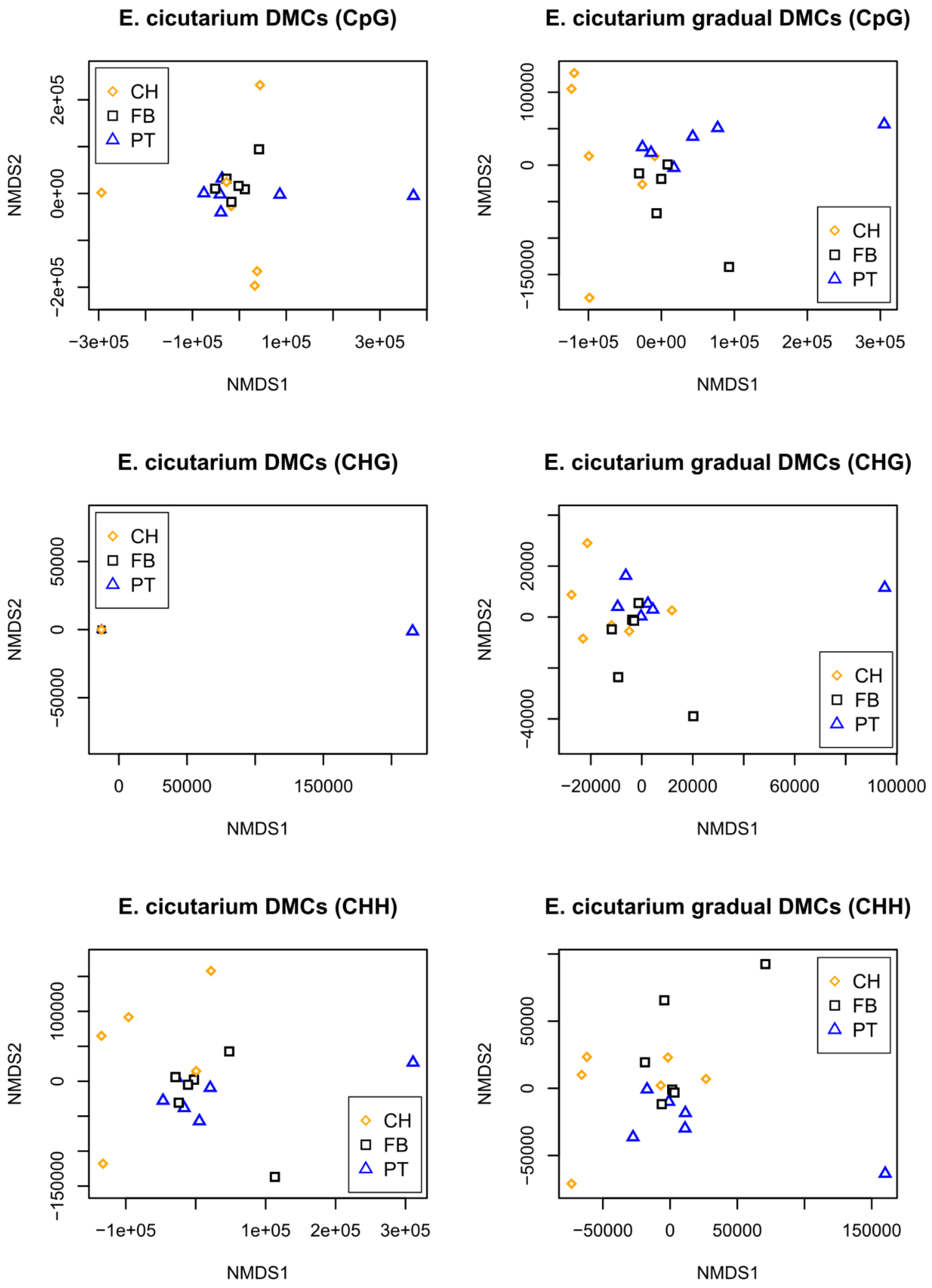


**Supplementary Figure S11.** Multilocus analysis of individual variation in methylation percentage of DMCs of *E. cicutarium* populations separated by context. CpG, CHG and CHH are shown from top to bottom rows, respectively. Left column shows the output of non-metric multidimensional scaling (NMDS) for all the DMCs, while the right column shows the output for the gradual DMCs. X and Y axes show NMDS analysis dimensions 1 and 2, respectively. Colors of data points show the population of origin (yellow: CH, black: FB, blue: PL).

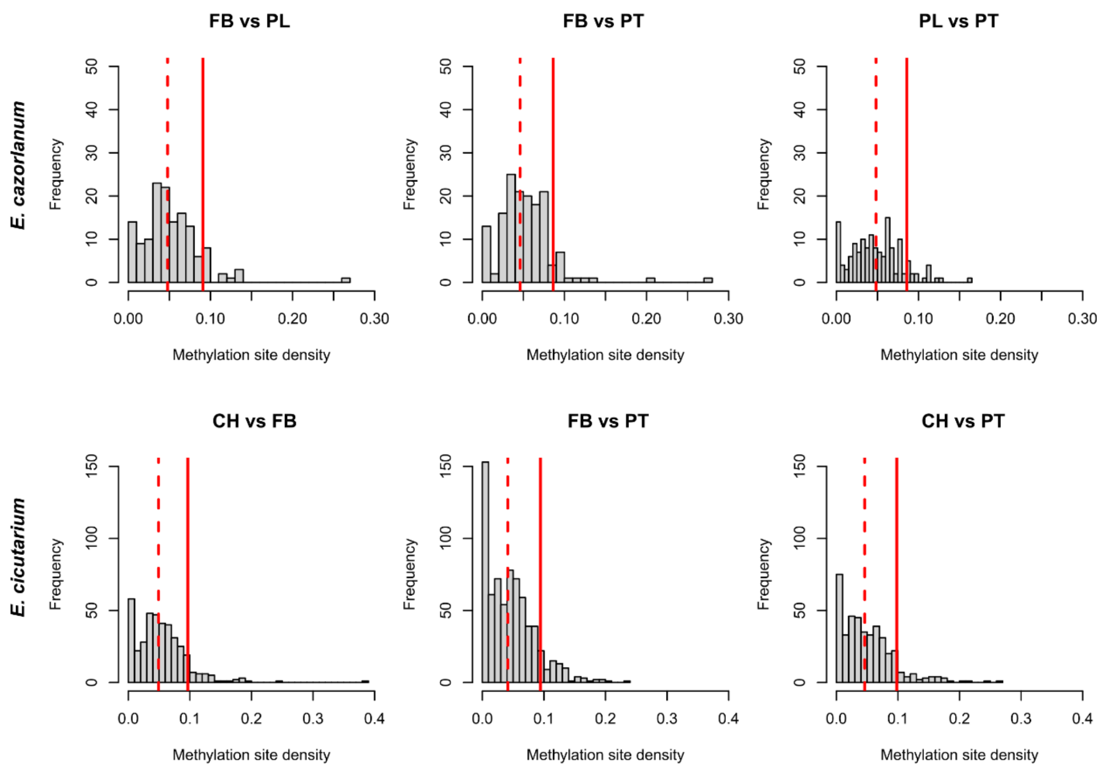


**Supplementary Figure S12.** Distribution of methylation site density (MSD) within DMRs defined between pairs of study populations of *E. cazorlanum* (upper row) and *E. cicutarium* (bottom row). Columns show population pairwise comparisons. X axis shows the distribution of MSD values (number of DMCs divided by the length of their respective DMR). Y axis shows the frequency of DMRs for each MSD specific value. Vertical red lines show the percentile 90 of each distribution used to define DMRs. Vertical dashed red lines show the median values of each distribution.

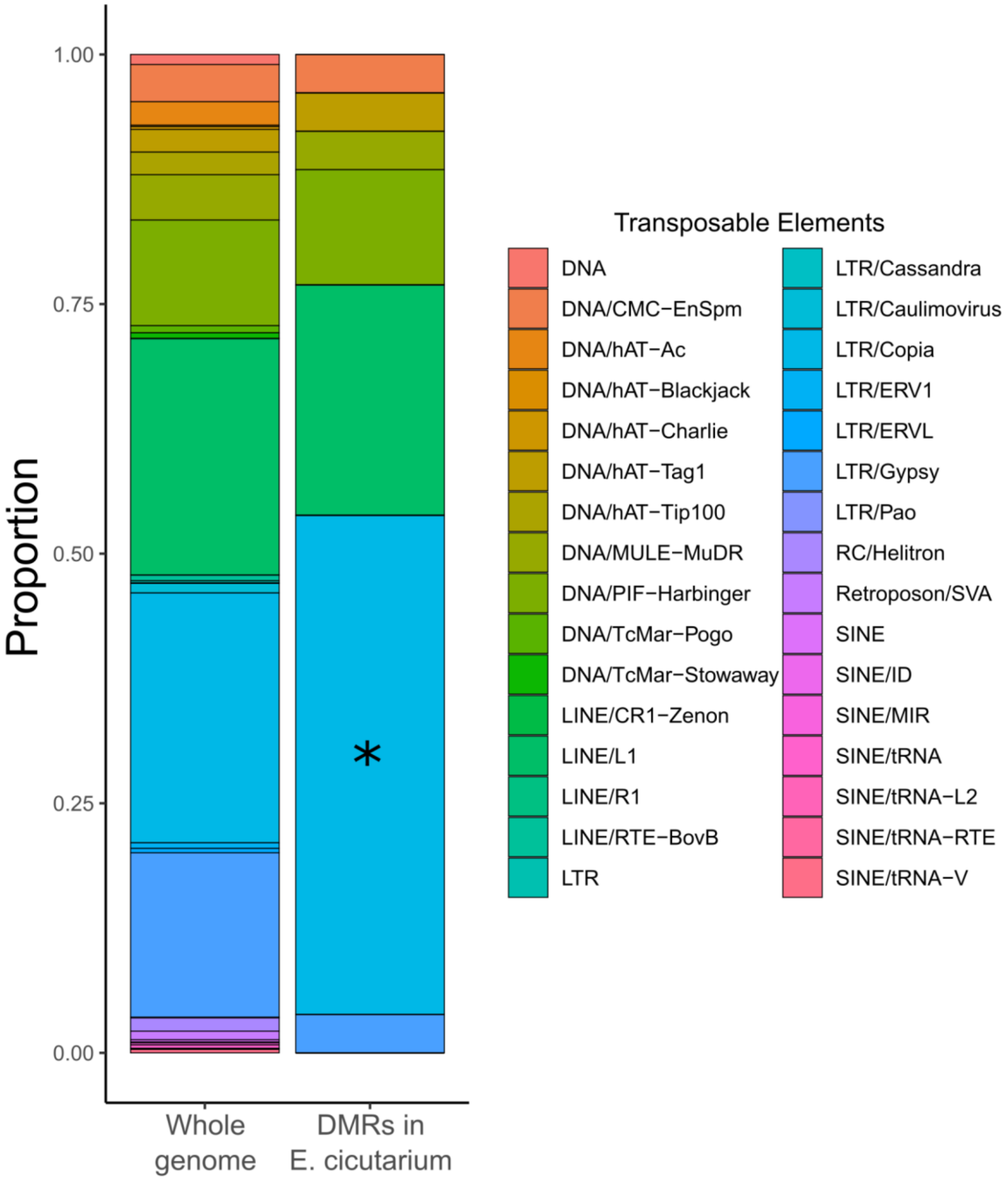


**Supplemental Figure S13.** Comparison between transposable elements (TEs) located in the *E. cicutarium* reference genome and those overlapping with differentially methylated regions. Y-axis shows the relative abundance (in proportion) of all repeated elements (including transposable elements) found in *E. cicutarium* genome (left bar, N = 397,130) or overlapping with a DMR found in the *E. cicutarium* dataset (right bar, N = 27). The asterisk indicates the statistical significance level of the Fisher's exact test for the TE group where it is placed (*: 0.05 > P-value > 0.01).

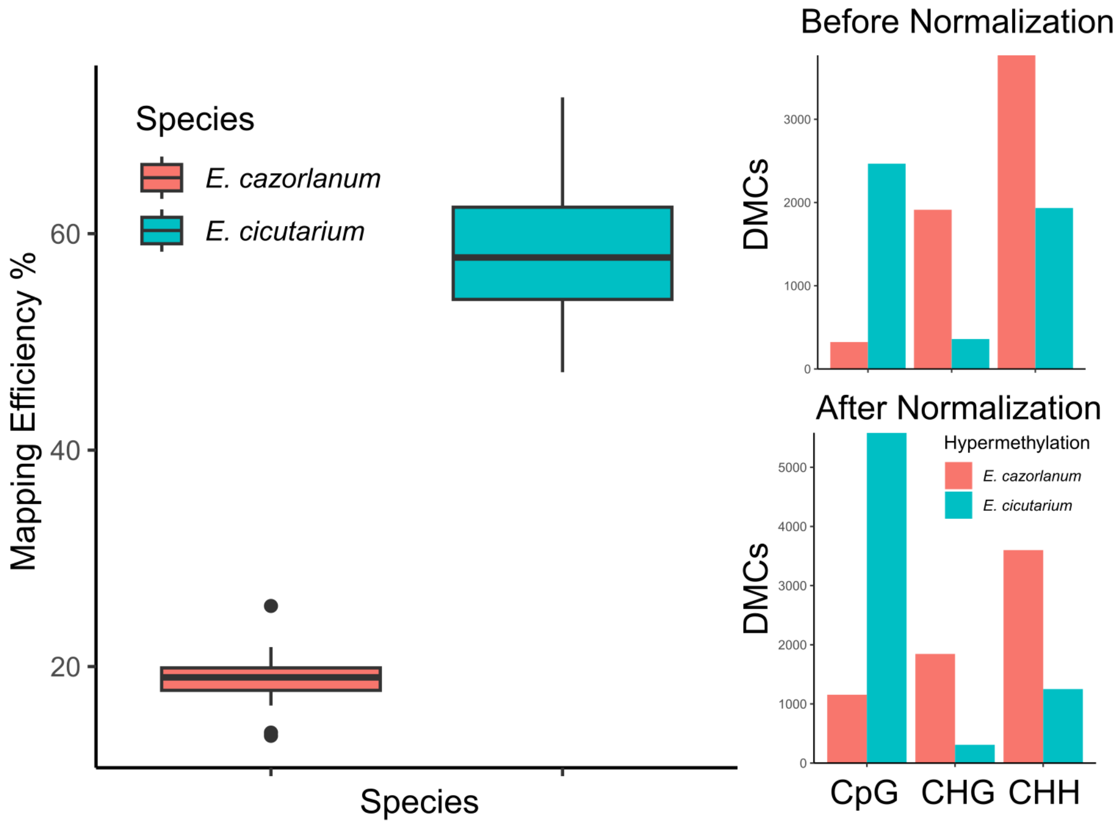


**Supplementary Figure S14.** Read alignment to draft genome efficiency and detection of DMCs based on bsRADseq. Differences in read alignment efficiency between *E. cazorlanum* and *E. cicutarium* bsRADseq reads are shown in the left plot. X axis shows the two *Erodium* species, Y axis shows the mapping efficiency percentage. Each box shows the median (central line), quartiles 1 and 3 (bottom and top edges of the box), maximum and minimum values (box whiskers) per species (represented by different colors). Right plots show the number of differentially methylated cytosines (DMCs) found in the inter-species comparison, before (top) and after (bottom) library normalization. Bars show hyper-methylated DMCs in *E. cazorlanum* (red) or *E. cicutarium* (blue). X-axis shows cytosine methylation contexts, Y-axis shows number of detected DMCs.

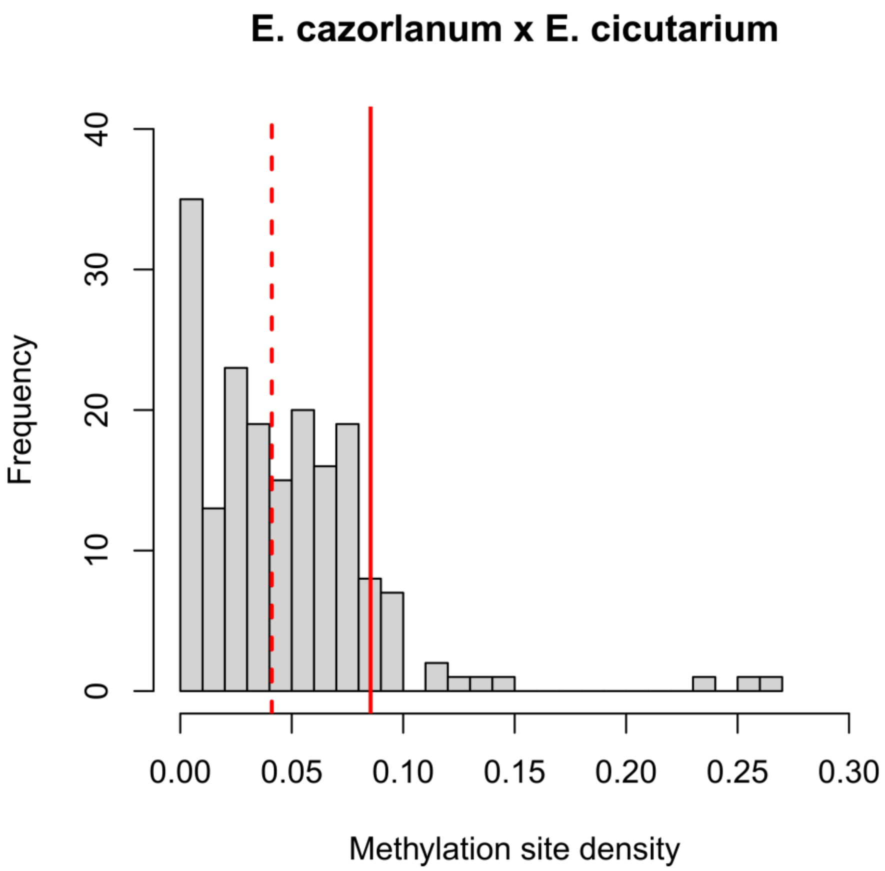


**Supplementary Figure S15.** Distribution of methylation site density (MSD) values per fragment for the comparison between the two studied *Erodium* species. X axis shows the MSD values (number of DMCs divided by the length of their respective DMR). Y-axis shows the frequency of DMRs for each MSD specific value. Vertical red line shows the percentile 90 of the distribution. Vertical dashed red line shows the median values of the distribution.

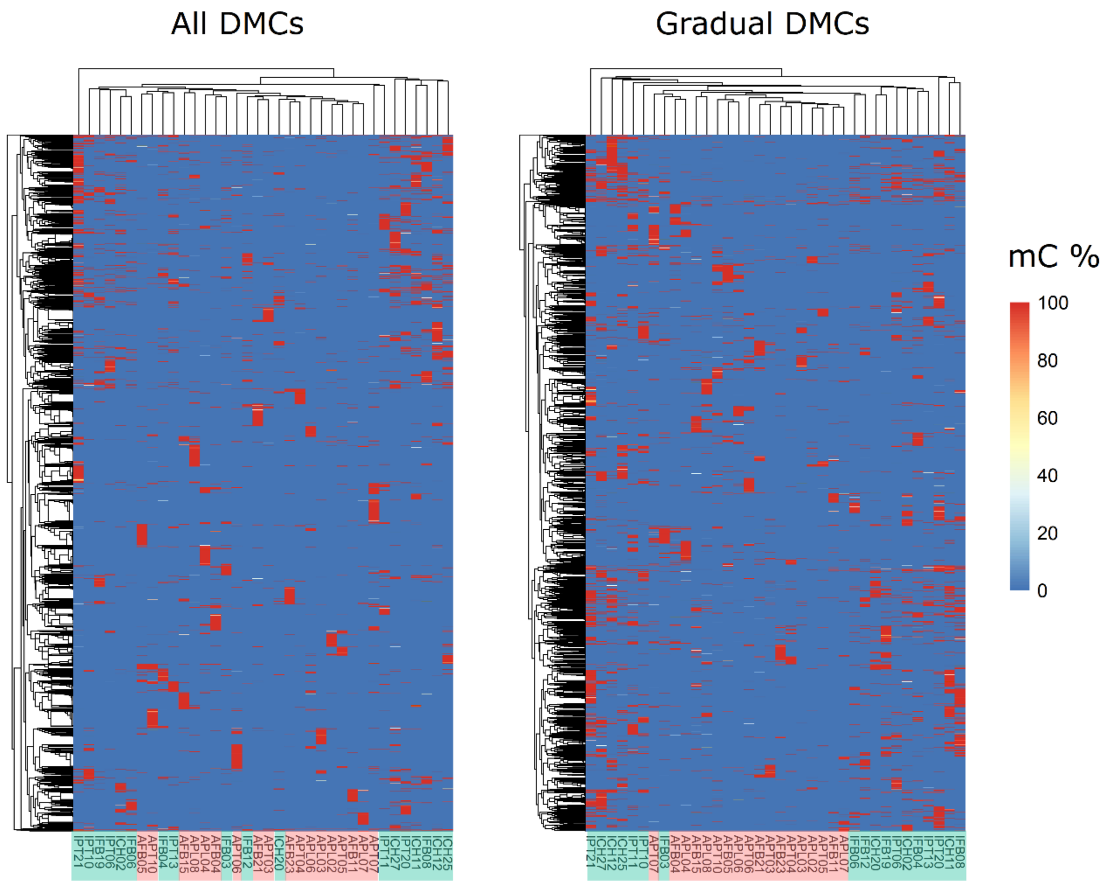


**Supplementary Figure S16.** Heatmap of DMC methylation percentage and hierarchical clustering of individuals based on percentage of methylation at DMCs detected in the inter-species comparison. X axis shows all individual samples arranged by hierarchical clustering. Y axis shows the different DMCs, arranged by hierarchical clustering. Colored tiles show the methylation percentage values per DMC and sample, ranging from 0 % (deep blue) to 100 % (red) methylation percentage. The left heatmap corresponds to the analysis including all DMCs, the right heatmap corresponds to the analysis with only gradual DMCs. Background of sample labels are colored based on the species (red: *E. cazorlanum*, green: *E. cicutarium*).

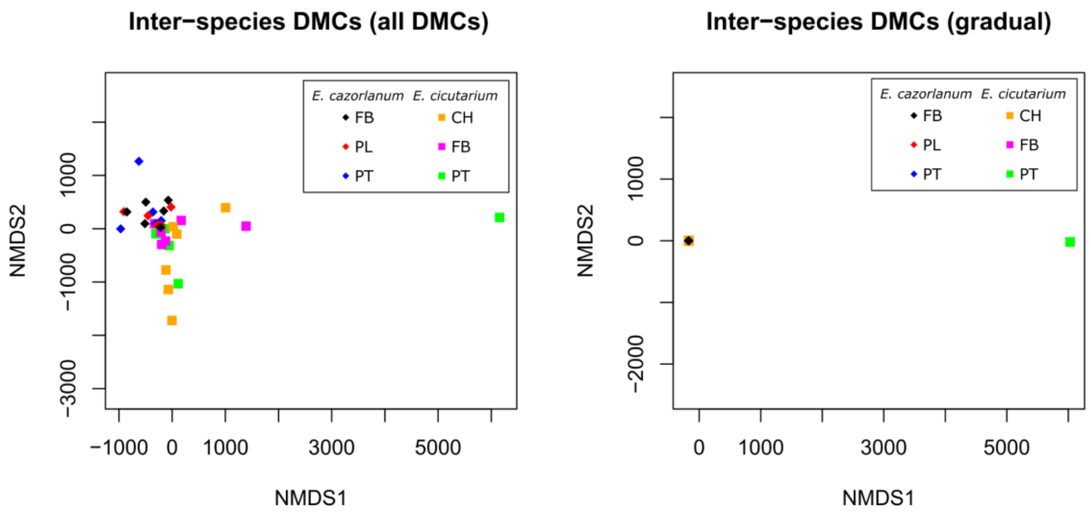


**Supplementary Figure S17.** Multilocus analysis of individual variation in methylation percentage of DMCs of *E. cazorlanum* and *E. cicutarium* populations in the inter-species comparison. Left column shows the output of non-metric multidimensional scaling (NMDS) for all the DMCs, while the right column shows the output for the gradual DMCs. X and Y axes show NMDS analysis dimensions 1 and 2, respectively. Colors of data points show the population of origin (black: FB, red: PL, blue: PT for *E. cazorlanum*; orange: CH, magenta: FB, green: PL for *E. cicutarium*).

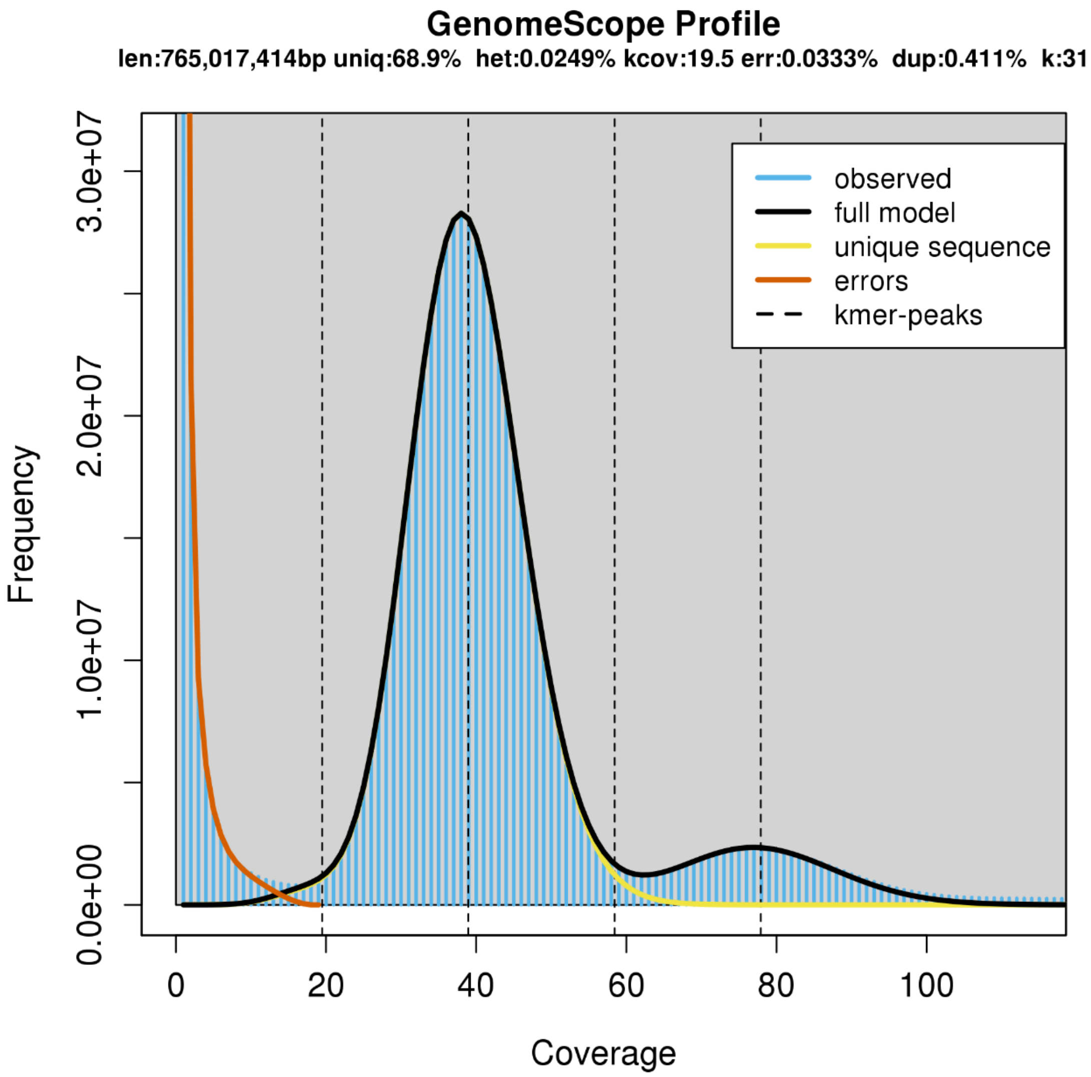


**Supplementary Figure S18.** GenomeScope K-mer = 31 graph, estimating coverage depth of reads for the genome of *Erodium cicutarium* using number of times a K-mer is observed (coverage) by number of K-mers with that coverage (frequency).

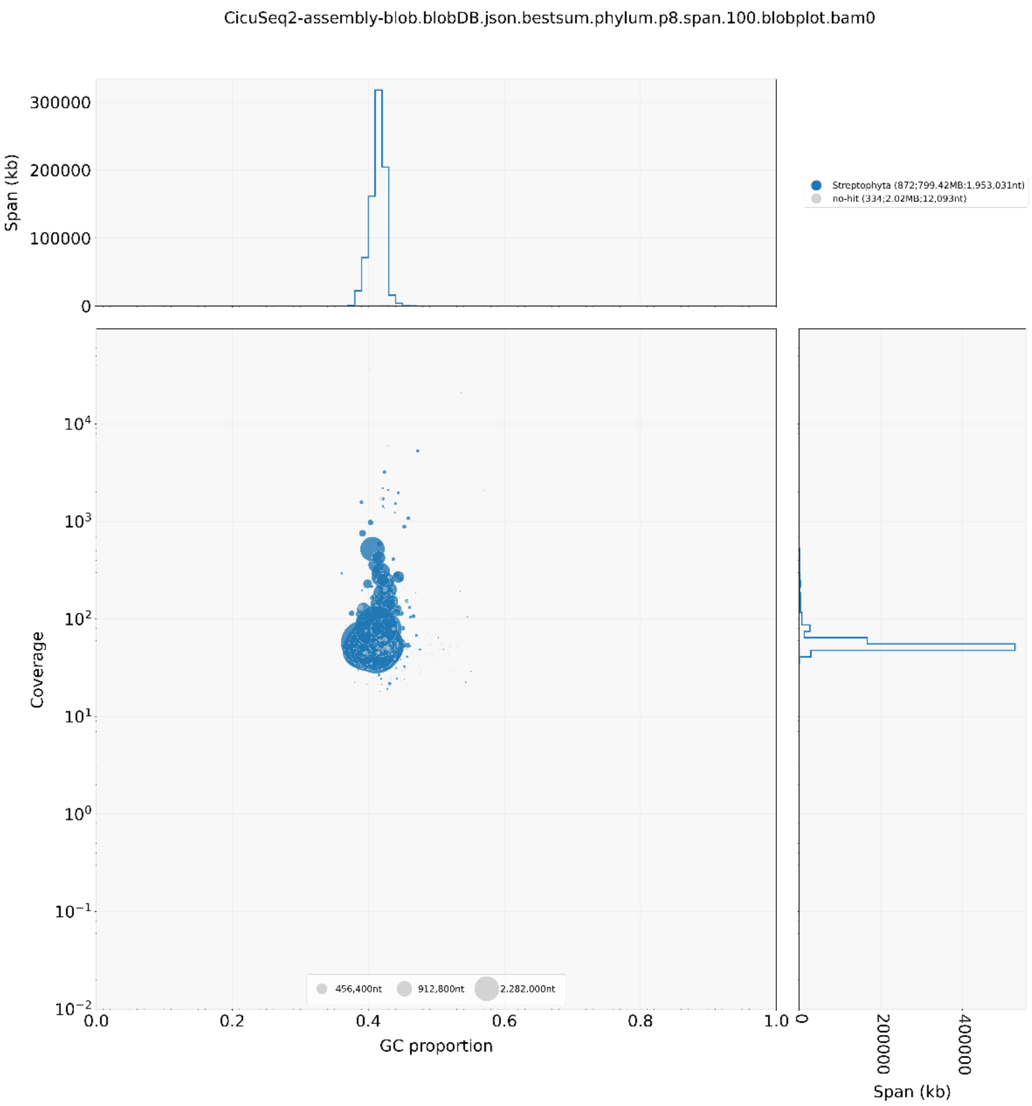


**Supplementary Figure S19.** Blobplot of the MaSuRCA polished assembly. The central plot shows the CG proportion (X-axis) compared to the coverage (Y-axis), circles represents the different contigs from the genome assembly, and their size represents the contig length. Color represents the annotation of each contig (blue: Streptophyta, grey: no-hit). The top and right plots show the distribution of genome sequence length covered per type of annotated subset (top: proportion of CG, right: coverage).

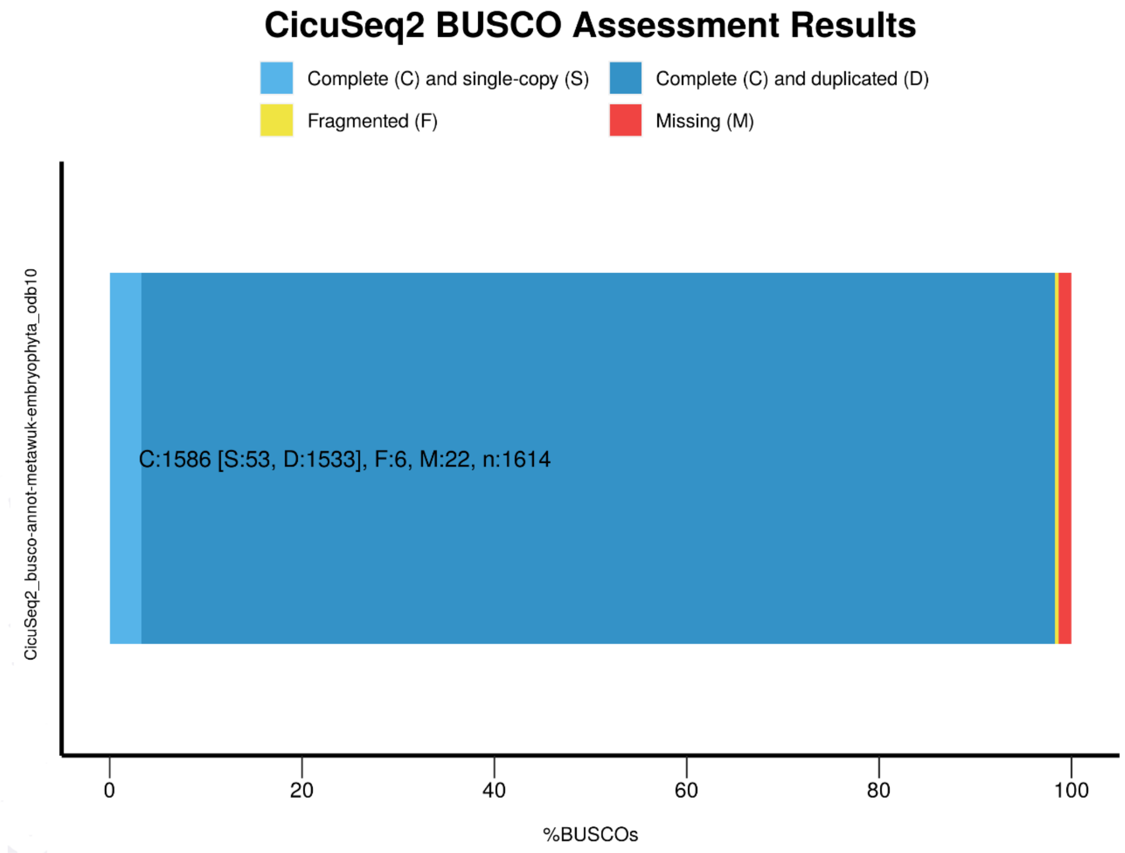


**Supplementary Figure S20.** Summary of BUSCO analysis results for the MaSurCa polished assembly. X-axis shows the percentage of BUSCOs found in the genome. Y-axis shows the analyzed set (in this case, the data comes from embryophyta obd10 database). Each color represents a different BUSCO category.

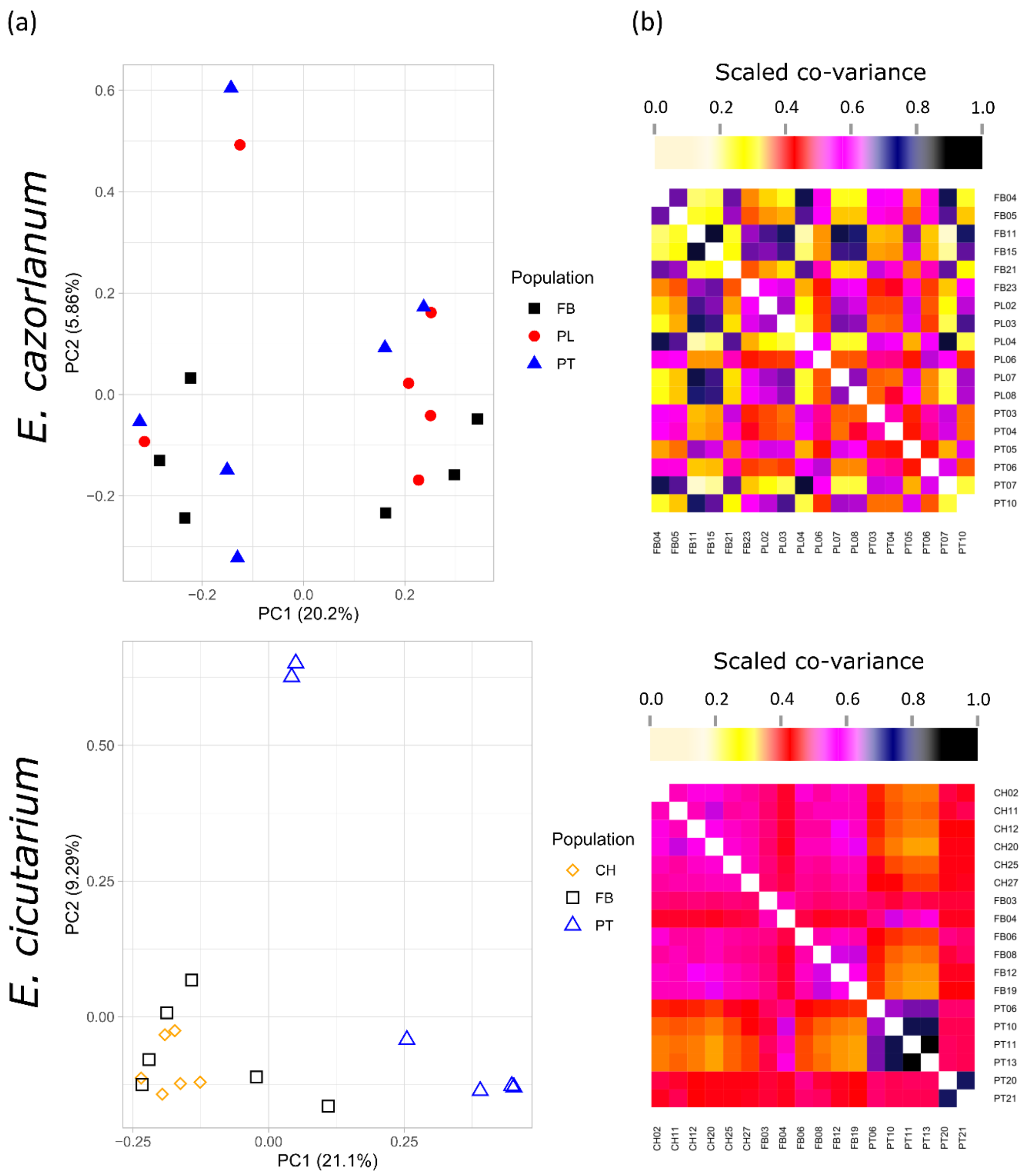


**Supplementary Figure S21.** Multivariate analysis of SNP calling. PCAs (a) and heatmaps of the co-variance matrix (b) from *E. cazorlanum* (top row) and *E. cicutarium* (bottom row). The PCA biplots show PCs 1 and 2 in X- and Y-axis, respectively; and the respective percentage of explained variance is shown between parentheses. Dots represent individual samples, and color and shape of the dot represent their source population. Heatmaps show the symmetric co-variance matrix with pair-wise comparisons; both axes show the sample labels sorted by population, and the intensity of the colour is proportional to the scaled co-variance.

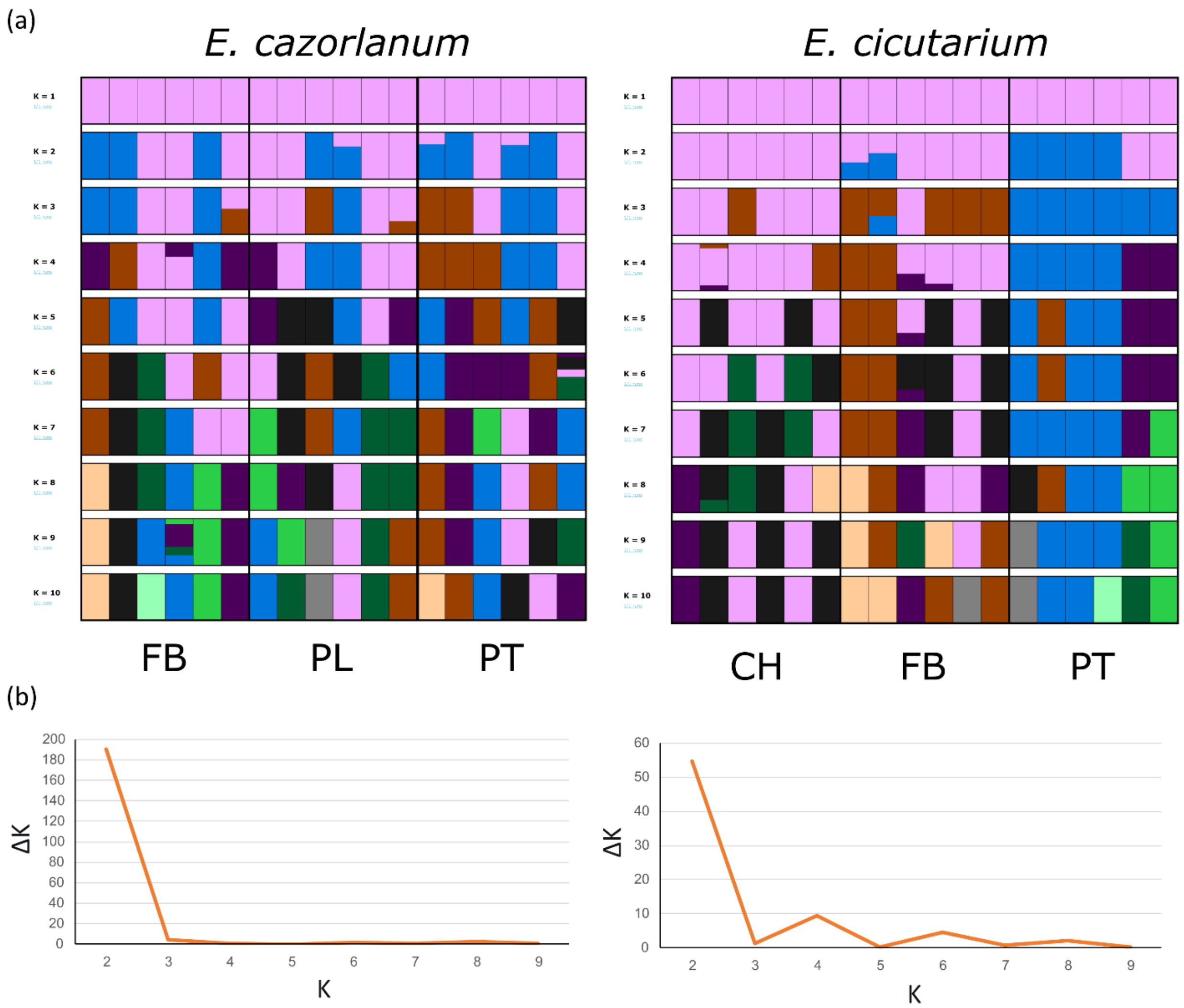


**Supplementary Figure S22.** Population genetic structure analysis. The top row shows admixture plots (a) and the bottom row shows the ΔK values from Evanno's method (b), for *E. cazorlanum* (left) and *E. cicutarium* (right), respectively. Each row from the admixture plots shows the admixture proportion per individual assuming a K value from 1 (top row) to 10 (bottom row), X-axis shows the admixture proportion per individual, Y-axis shows all study individuals arranged by population of origin, boxes with thicker lines determine the three different populations, and different colors show a different ancestry.

#### References

Akiva, E., Brown, S., Almonacid, D. E., Barber, A. E., 2nd, Custer, A. F., Hicks, M. A., Huang, C. C., Lauck, F., Mashiyama, S. T., Meng, E. C., Mischel, D., Morris, J. H., Ojha, S., Schnoes, A. M., Stryke, D., Yunes, J., M.Ferrin, T. E., Holliday, G. L., & Babbitt, P. C. (2014). The Structure-Function Linkage Database. *Nucleic Acids Research,* **42**(Database issue), D521-530. doi:10.1093/nar/gkt1130

Andrews, S. R. (2010). FastQC: a quality control tool for high throughput sequence data. Retrieved from protocols.io

Attwood, T. K., Beck, M. E., Bleasby, A. J., & Parry-Smith, D. J. (1994). PRINTS a database of protein motif fingerprints. *Nucleic Acids Research,* **22**(17), 3590-3596.

Behr, A. A., Liu, K. Z., Liu-Fang, G., Nakka, P., & Ramachandran, S. (2016). pong: fast analysis and visualization of latent clusters in population genetic data. *Bioinformatics,* **32**(18), 2817-2823. doi:10.1093/bioinformatics/btw327

Consortium, U. (2007). The universal protein resource (UniProt). *Nucleic Acids Research,* **36**(suppl_1), D190-D195.

Evanno, G., Regnaut, S., & Goudet, J. (2005). Detecting the number of clusters of individuals using the software STRUCTURE: a simulation study. *Molecular Ecology,* **14**(8), 2611-2620. doi:10.1111/j.1365-294X.2005.02553.x

Fadiji, A. E., & Babalola, O. O. (2020). Metagenomics methods for the study of plant-associated microbial communities: A review. *Journal of Microbiology Methods,* **170**, 105860. doi:10.1016/j.mimet.2020.105860

Finn, R. D., Bateman, A., Clements, J., Coggill, P., Eberhardt, R. Y., Eddy, S. R., Heger, A., Hetherington, K., Holm, L., Mistry, J., Sonnhammer, E. L., Tate, J., & Punta,

M. (2014). Pfam: the protein families database. *Nucleic Acids Research,* **42**(Database issue), D222-230. doi:10.1093/nar/gkt1223

Flynn, J. M., Hubley, R., Goubert, C., Rosen, J., Clark, A. G., Feschotte, C., & Smit, A. F. (2020). RepeatModeler2 for automated genomic discovery of transposable element families. *Procceedings of the National Academy of Science of the USA,* **117**(17), 9451-9457. doi:10.1073/pnas.1921046117

Gough, J., Karplus, K., Hughey, R., & Chothia, C. (2001). Assignment of homology to genome sequences using a library of hidden Markov models that represent all proteins of known structure. *Journal of Molecular Biology,* **313**(4), 903-919. doi:10.1006/jmbi.2001.5080

Gurevich, A., Saveliev, V., Vyahhi, N., & Tesler, G. (2013). QUAST: quality assessment tool for genome assemblies. *Bioinformatics,* **29**(8), 1072-1075. doi:10.1093/bioinformatics/btt086

Haft, D. H., Selengut, J. D., Richter, R. A., Harkins, D., Basu, M. K., & Beck, E. (2013). TIGRFAMs and Genome Properties in 2013. *Nucleic Acids Research,* **41**(Database issue), D387-395. doi:10.1093/nar/gks1234

Haghshenas, E., Asghari, H., Stoye, J., Chauve, C., & Hach, F. (2020). HASLR: Fast Hybrid Assembly of Long Reads. *iScience,* **23**(8), 101389. doi:10.1016/j.isci.2020.101389

Hufnagel, D. E., Hufford, M. B., & Seetharam, A. S. (2020). SequelTools: a suite of tools for working with PacBio Sequel raw sequence data. *BMC Bioinformatics,* **21**(1), 429. doi:10.1186/s12859-020-03751-8

Jones, P., Binns, D., Chang, H. Y., Fraser, M., Li, W., McAnulla, C., McWilliam, H., Maslen, J., Mitchell, A., Nuka, G., Pesseat, S., Quinn, A. F., Sangrador-Vegas, A., Scheremetjew, M., Yong, S. Y., Lopez, R., & Hunter, S. (2014). InterProScan

5: genome-scale protein function classification. *Bioinformatics,* **30**(9), 1236-1240. doi:10.1093/bioinformatics/btu031

Kollmar, M. (2019). *Gene Prediction: Methods and Protocols* (Vol. 1962): Springer Nature.

Kolmogorov, M., Yuan, J., Lin, Y., & Pevzner, P. A. (2019). Assembly of long, error-prone reads using repeat graphs. *Nature Biotechnology,* **37**(5), 540-546. doi:10.1038/s41587-019-0072-8

Korneliussen, T. S., Albrechtsen, A., & Nielsen, R. (2014). ANGSD: Analysis of Next Generation Sequencing Data. *BMC Bioinformatics,* **15**(356). doi: 10.1186/s12859-014-0356-4

Langmead, B., & Salzberg, S. L. (2012). Fast gapped-read alignment with Bowtie 2. *Nature Methods,* **9**(4), 357-359. doi:10.1038/nmeth.1923

Letunic, I., & Bork, P. (2018). 20 years of the SMART protein domain annotation resource. *Nucleic Acids Research,* **46**(D1), D493-D496. doi:10.1093/nar/gkx922

Lewis, T. E., Sillitoe, I., Dawson, N., Lam, S. D., Clarke, T., Lee, D., Orengo, C., & Lees, J. (2018). Gene3D: Extensive prediction of globular domains in proteins. *Nucleic Acids Research,* **46**(D1), D435-D439. doi:10.1093/nar/gkx1069

Li, H. (2013). Aligning sequence reads, clone sequences and assembly contigs with BWA-MEM. *arXiv*(1-3).

Li, H., Handsaker, B., Wysoker, A., Fennell, T., Ruan, J., Homer, N., Marth, G., Abecasis, G., Durbin, R., & Subgroup of Genome Project Data Processing. (2009). The Sequence Alignment/Map format and SAMtools. *Bioinformatics,* **25**(16), 2078-2079. doi:10.1093/bioinformatics/btp352

Lu, S., Wang, J., Chitsaz, F., Derbyshire, M. K., Geer, R. C., Gonzales, N. R., Gwadz, M., Hurwitz, D. I., Marchler, G. H., Song, J. S., Thanki, N., Yamashita, R. A.,

Yang, M., Zhang, D., Zheng, C., Lanczycki, C. J., & Marchler-Bauer, A. (2020). CDD/SPARCLE: the conserved domain database in 2020. *Nucleic Acids Research,* **48**(D1), D265-D268. doi:10.1093/nar/gkz991

Lupas, A., Van Dyke, M., & Stock, J. (1991). Predicting Coiled Coils from Protein Sequences. *Science,* **252**(5009), 1162-1164.

Marcais, G., & Kingsford, C. (2011). A fast, lock-free approach for efficient parallel counting of occurrences of k-mers. *Bioinformatics,* **27**(6), 764-770. doi:10.1093/bioinformatics/btr011

Marcais, G., Yorke, J. A., & Zimin, A. (2015). QuorUM: An Error Corrector for Illumina Reads. *PLoS One,* **10**(6), e0130821. doi:10.1371/journal.pone.0130821

Meisner, J., & Albrechtsen, A. (2018). Inferring Population Structure and Admixture Proportions in Low-Depth NGS Data. *Genetics,* **210**(2), 719-731. doi:10.1534/genetics.118.301336

Mi, H., Muruganujan, A., & Thomas, P. D. (2013). PANTHER in 2013: modeling the evolution of gene function, and other gene attributes, in the context of phylogenetic trees. *Nucleic Acids Research,* **41**(Database issue), D377-386. doi:10.1093/nar/gks1118

Munoz-Merida, A., Viguera, E., Claros, M. G., Trelles, O., & Perez-Pulido, A. J. (2014). Sma3s: a three-step modular annotator for large sequence datasets. *DNA Research,* **21**(4), 341-353. doi:10.1093/dnares/dsu001

Necci, M., Piovesan, D., Dosztanyi, Z., & Tosatto, S. C. E. (2017). MobiDB-lite: fast and highly specific consensus prediction of intrinsic disorder in proteins. *Bioinformatics,* **33**(9), 1402-1404. doi:10.1093/bioinformatics/btx015

Oksanen, J., Simpson, G. L., Blanchet, F. G., Kindt, R., Legendre, P., Minchin, P. R., O'Hara,R.B., Solymos, P., Stevens, M.H.H., Szoecs, E., Wagner, H., Barbour, M.,

Bedward, M., Bolker, B., Borcard, D., Carvalho, G., Chirico, M., De Caceres, M., Durand, S., Evangelista, H.B.A., FitzJohn, R., Friendly, M., Furneaux, B., Hannigan, G., Hill, M.O., Lahti, L., McGlinn, D., Ouellette, M-H., Cunha, E.R., Smith, T., Stier, A., Ter Braak, C.J.F., 6 Weedon, J. (2022). vegan: Community Ecology Package. Retrieved from https://CRAN.R-project.org/package=vegan

Pedruzzi, I., Rivoire, C., Auchincloss, A. H., Coudert, E., Keller, G., de Castro, E., Baratin, D., Cuche, B. A., Bougueleret, L., Poux, S., Redaschi, N., Xenarios, I., & Bridge, A. (2015). HAMAP in 2015: updates to the protein family classification and annotation system. *Nucleic Acids Res,* **43**(Database issue), D1064-1070. doi:10.1093/nar/gku1002

Pertea, G., & Pertea, M. (2020). GFF Utilities: GffRead and GffCompare. *F1000Res, 9*. doi:10.12688/f1000research.23297.2

Pritchard, J. K., Stephens, M., & Donnelly, P. (2000). Inference of Population Structure Using Multilocus Genotype Data. *Genetics,* **155**(2), 945-959. Doi: 10.1093/genetics/155.2.945

Schalamun, M., & Schwessinger, B. ( 2017). DNA size selection (>1kb) and clean up using an optimized SPRI beads mixture. Retrieved from  protocols.io.

Sigrist, C. J., de Castro, E., Cerutti, L., Cuche, B. A., Hulo, N., Bridge, A., Bougueleret, L., & Xenarios, I. (2013). New and continuing developments at PROSITE. *Nucleic Acids Research,* **41**(Database issue), D344-347. doi:10.1093/nar/gks1067

Skotte, L., Korneliussen, T. S., & Albrechtsen, A. (2013). Estimating individual admixture proportions from next generation sequencing data. *Genetics,* **195**(3), 693-702. doi:10.1534/genetics.113.154138

Smit, A., Hubley, R., & Green, P. (2021). 2013–2015. RepeatMasker Open-4.0. https://www.repeatmasker.org/

Stanke, M., Keller, O., Gunduz, I., Hayes, A., Waack, S., & Morgenstern, B. (2006). AUGUSTUS: ab initio prediction of alternative transcripts. *Nucleic Acids Research,* **34**(Web Server issue), W435-439. doi:10.1093/nar/gkl200

R Core Team (2023). R: A Language and Environment for Statistical Computing. Retrieved from https://www.R-project.org/

Vurture, G. W., Sedlazeck, F. J., Nattestad, M., Underwood, C. J., Fang, H., Gurtowski, J., & Schatz, M. C. (2017). GenomeScope: fast reference-free genome profiling from short reads. *Bioinformatics,* **33**(14), 2202-2204. doi:10.1093/bioinformatics/btx153

Wu, C. H., Nikolskaya, A., Huang, H., Yeh, L. S., Natale, D. A., Vinayaka, C. R., Hu, Z. Z., Mazumder, R, .Kumar, S., Kourtesis, P., Ledley, R. S., Suzek, B. E., Arminski, L., Chen, Y., Zhang, J., Cardenas, J. L., Chung, S., Castro-Alvear, J., Dinkov, G., & Barker, W. C. (2004). PIRSF: family classification system at the Protein Information Resource. *Nucleic Acids Research,* **32**(Database issue), D112-114. doi:10.1093/nar/gkh097

Zimin, A. V., Marcais, G., Puiu, D., Roberts, M., Salzberg, S. L., & Yorke, J. A. (2013). The MaSuRCA genome assembler. *Bioinformatics,* **29**(21), 2669-2677. doi:10.1093/bioinformatics/btt476

Zimin, A. V., Puiu, D., Luo, M. C., Zhu, T., Koren, S., Marcais, G., Yorke, J. A. Dvorak, J., & Salzberg, S. L. (2017). Hybrid assembly of the large and highly repetitive genome of Aegilops tauschii, a progenitor of bread wheat, with the MaSuRCA mega-reads algorithm. *Genome Research,* **27**(5), 787-792. doi:10.1101/gr.213405.116

Zimin, A. V., & Salzberg, S. L. (2020). The genome polishing tool POLCA makes fast and accurate corrections in genome assemblies. *PLoS Computational Biology,* **16**(6), e1007981. doi:10.1371/journal.pcbi.1007981
